# Two genomes in one mitochondrion: mitochondrial heteroplasmy and mitogenome dynamics in *Chlamydomonas reinhardtii*

**DOI:** 10.64898/2026.10.29.759696

**Authors:** Zhongshi He, Tsvetan R. Bachvaroff, Sean D. Gallaher, Yantao Li

## Abstract

The mitochondrial genome of the model green alga *Chlamydomonas reinhardtii* has long been described as a 15.8-kb linear molecule capped by inverted telomeres. Leveraging high-accuracy long-read sequencing to overcome limitations associated with conventional genome algorithm biases, we demonstrated that this classical paradigm was incomplete. Our analyses provided no evidence for canonical telomeric structures, instead revealed widespread structural heteroplasmy characterized by the coexistence of two distinct mitochondrial genome configurations. In addition to the historical 15.8-kb configuration (designated as config 2), an expanded 19-kb configuration (config 1) was found containing 4,001-bp terminal inverted repeats that duplicate the *cob* and *nad4* genes. Single-molecule long-read profiling provided many reads longer than a single config 1 or 2 monomer and captured a dynamic network of multimeric mitochondrial DNA concatemers bridged with a conserved 17-bp homologous motif. Together, these structural features support a recombination-dependent replication (RDR) model that could enable mitochondrial DNA replication in the absence of conventional telomeric structures and reverse transcriptase-like (*rtl*) genes. We further identify widespread nuclear mitochondrial DNA segments (NUMTs), which complicate targeted amplification and quantification of these mitogenome configurations. Specifying NUMTs enabled configuration-specific digital PCR measurements across the laboratory lineage and additional field-isolated *Chlamydomonas* strains, revealing substantial variation in the relative abundance of the two mitogenome configurations. These findings revise a four-decade-old model of the *C. reinhardtii* mitogenome and provided a new framework for investigating mitochondrial genome architecture, structural heteroplasmy, and replication dynamics in photosynthetic eukaryotes.

**Significance statement:** For over four decades, the mitochondrial genome of the model alga *Chlamydomonas reinhardtii* has been believed to be a streamlined, linear molecule capped by telomeres. Here, we revised this long-standing paradigm by showing that the mitogenome lacked canonical telomeric structures previously proposed. Instead, it exhibited widespread structural heteroplasmy characterized by the coexistence of two distinct mitogenome configurations and dynamic multimeric concatemers. These structural features supported a recombination-dependent replication model. We further identified widespread nuclear mitochondrial DNA segments (NUMTs) that confounded mitogenome-specific amplification and quantification. By resolving these two configurations and quantify their absolute abundances across laboratory and field-isolated strains, our findings provide a new framework for studying organelle genome architecture, replication mechanisms, and evolution in photosynthetic eukaryotes.

## Introduction

As a unicellular model organism, the green alga *Chlamydomonas reinhardtii* has been extensively utilized across diverse areas of biology, including cellular metabolism, organelle biology, functional genetics, and synthetic biology (1). Its mitochondrion has also been extensively studied at the genomic to proteomic level (2), providing an important model for investigating mitochondrial biology and methodological development in photosynthetic eukaryotes. Early electron microscopy revealed that the *C. reinhardtii* mitogenome was predominantly linear, with a rare fraction (≤ 1%) occurring as open and supercoiled circular molecules ranging from 4.0 to 5.4 µm in length (3). Subsequent work by Boer and Gray established a size of 15.8 kb for this linear genome and identified its core components (4), including the *nad1* and *rtl* genes (5), three tRNAs (6), and fragmented large and small subunit (LSU and SSU) rRNAs (7). A complete genetic map was subsequently generated through Sanger sequencing by Michaelis et al. (8) and later refined using Illumina short-read sequencing by Gallaher et al. (9). The resulting canonical mitogenome was remarkably streamlined and included seven respiratory chain protein-coding genes (*nad1*, *nad2*, *nad4*, *nad5*, *nad6*, *cob*, and *cox1*), 14 discontinuous rRNA fragments (*rrnL6*, *S1*, *L5*, *L7*, *S2*, *L3a*, *L3b*, *L8*, *S3*, *L4*, *S4*, *L1*, *L2a*, and *L2b*), three tRNAs (trnW-CCA, trnQ-UUG, and trnM-CAU), and an open reading frame encoding a reverse transcriptase-like protein (*rtl*) (9).

Specific structural features of the *C. reinhardtii* mitogenome provided the basis for two proposed mitochondrial DNA replication models. The first, proposed by Vahrenholz and colleagues, relied on an internal 86-nucleotide repeat that perfectly matches the terminal sequences of the linear mitogenome or so called “telomeres,” including their single-stranded 3’ extensions (10). In this model, transcription of the internal repeat generates an RNA primer that initiates DNA synthesis at the genome termini, driving a dual leading-strand replication. Subsequently, the *rtl*-encoded reverse transcriptase facilitates terminal completion by utilizing parental strand-associated RNA primers. This RNA-primed model provided a potential explanation for the maintenance of the linear telomeres, the internal 86-bp repeat, and the conserved *rtl* gene. An alternative DNA-based model proposed that intramolecular folding aligns the terminal sequence and internal repeats to initiate DNA synthesis (11). In this T-loop model, the left telomere folds back and invades the homologous internal repeat, generating a circular intermediate while enabling the 3’ telomeric overhang to prime DNA synthesis. Because these models elegantly provided plausible mechanisms for overcoming the end-replication problems inherent to a linear mitogenome (12) and explained the presence of the conserved *rtl* gene, they have been wildly accepted for decades. However, both models rely on definite sequence architectures at the mitogenome termini (whether identical internal repeats or 3’ telomeric overhangs), whereas these regions are structurally fluid across species and notoriously difficult to resolve with short-reads sequencing (13).

Mitochondrial heteroplasmy is defined as the coexistence of distinct mitochondrial DNA (mtDNA) variants within a single cell (14). Although mitochondrial genomes in plants and algae are known to be uniparentally inherited (15), specific genetic or environmental conditions occasionally facilitate the coexistence of divergent mitogenomes (16). This heteroplasmic state—whether introduced by occasional paternal leakage and *de novo* mutations, or amplified through substoichiometric shifting—is often modulated by vegetative segregation and has been documented in wild carrot (*Daucus carota*) (17), bladder campion (*Silene vulgaris*) (18), and Stylonematophyceae red alga *Rhodosorus marinus* (19). Prior to the advent of high-throughput sequencing, electron microscopy provided structural evidence of this mitochondrial heterogeneity in plant cells (20). Rather than the expected circular genome structures, early ultrastructural analyses revealed complex assemblies of branched linear molecules, simple linear fragments, and subgenome-sized minicircles (21, 22). This structural and genetic complexity provides templates and opportunities for homologous recombination among distinct mitochondrial DNA molecules (23). However, traditional short-read sequencing and targeted quantitative assays struggle to fully capture this structural dynamism, as they cannot span complex recombination junctions. Addressing these limitations requires advanced methodologies capable of either *in vivo* monitoring or complete structural assembly. While real-time single-cell tracking provides a framework for observing variant segregation in yeast (24), single-molecule long-read sequencing offers the definitive solution for genomic architecture. This approach enables the direct resolution of native, heteroplasmic mitogenomes and the precise mapping of structural variants.

Our recent whole-genome sequencing of *C. reinhardtii* CC-5325 yielded a significant number of high-accuracy long reads, providing substantially improved resolution of *Chlamydomonas* mitogenome structure. We hypothesized that the established canonical linear map might not fully represent the *in vivo* structural heterogeneity of the mitogenome, thereby limiting our understanding of its replication and maintenance. To test this hypothesis, we developed new bioinformatic approaches to reassemble and reannotate the CC-5325 mitogenome and systematically examined the organization of its coding sequences, noncoding intergenic spacers, and terminal regions. By directly resolving alternative mitochondrial DNA configurations from individual long reads, we revealed previously unrecognized structural heteroplasmy with two mitogenome configurations. We also identified widespread nuclear mitochondrial DNA segments (NUMTs), which enabled configuration-specific quantification across the CC-5325 lineage and other field-isolated strains. Furthermore, these structural features challenge the previously proposed terminal architecture of the mitogenome and support a recombination-dependent replication (RDR) model of *Chlamydomonas* mitogenome replication.

## Results

### Long-read sequencing reveals alternative mitogenome configurations in *C. reinhardtii*

We isolated organelle-derived sequence from CC-5325 PacBio HiFi and Oxford Nanopore Technology (ONT) reads by mapping the raw datasets to the CC-503 plastome and mitogenome references (9). This enrichment yielded 160,467 high-quality organellar reads with an N50 of 15,543 bp, comparable to the length of previous published *C. reinhardtii* mitogenome assemblies. Subsequent *de novo* assembly of these reads produced a 19,039-bp contig, approximately 3.2 kb longer than the canonical ∼15.8-kb reference (Table 1). While annotation revealed this expanded mitogenome retained all expected functional features, it contained an extended terminal region with *cob*, *nad4*, and a partial *nad5* gene on the opposite strand, forming a 4,001-bp inverted repeat with the 5′ end of the genome (Fig. 1*A*).

**Fig. 1.**
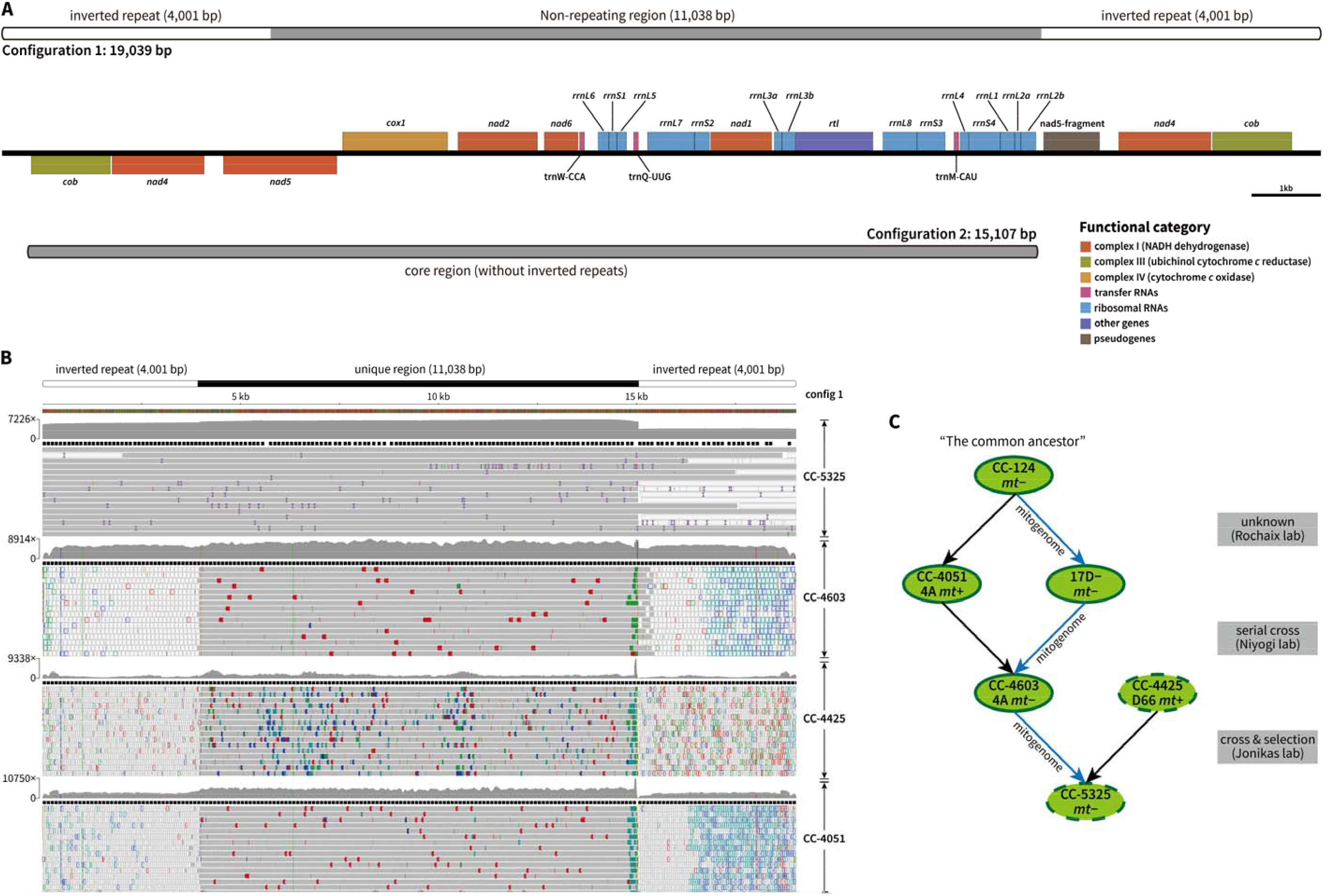
Structural characterization and lineage inheritance of two mitogenome configurations in *C. reinhardtii*. (A) Mitogenome structures of config 1 (19,039 bp) and config 2 (15,107 bp) assembled from long-read sequencing data. Boxes denote the locations of annotated mitochondrial genes, color-coded by their functional categories. (B) WGS read-alignment profiles of selected strains within the CC-5325 lineage, mapped to the config 1 mitogenome, and visualized using IGV. Gray tracks indicate high-confidence alignments with a mapping quality (MAPQ) score of 60; colored indicators denote structural variants or sequence mismatches of individual reads according to the IGV alignment interpretation guidelines (https://igv.org/doc/desktop/#UserGuide/tracks/alignments/). White tracks represent multi-mapping reads (MAPQ = 0) associated with the 4,001-bp terminal inverted repeats. (C) Genetic relationships among strains of the CC-5325 lineage. Blue arrows indicate the direction of predominantly uniparental mitogenome inheritance from the *mt^−^* parent through successive crosses and strain-selections events.

**Table 1:** Comparison of published mitogenomic sequences of *Chlamydomonas reinhardtii*.

| Strains | CCAP 11/32 *mt*<sup>+</sup>* | CW15 *mt*<sup>+</sup>* | CC-124 *mt*<sup>-</sup> | CC-503 *mt*<sup>-</sup> | CC-4532 *mt*<sup>-</sup> | CC-5325 *mt*<sup>-</sup>** |
| --- | --- | --- | --- | --- | --- | --- |
| *Genome information* |  |  |  |  |  |  |
| Published date | 1990* | 1993* | 2008 | 2018 | 2022 | 2026 |
| Provided database | GenBank | GenBank | GenBank | Literature | Phytozome | This study |
| Accession number | NC_001638.1 | U03843.1 | EU306622.1 | (9) | genome ID: 707 | - |
| Genome length (bp) | 15,758 | 15,758 | 15,787 | 15,789 | 15,789 | 19,039 |
| GC contents (%) | 45.20 | 45.20 | 45.25 | 45.24 | 45.24 | 44.98 |
| *Variant information* ** |  |  |  |  |  |  |
| SNPs | 23 | 23 | 10 | 9 | 9 | Query Baseline |
| single-base INDELs | 33 | 33 | 0 | 0 | 0 |  |
| Insertion count (>1 bp) | 3 | 3 | 3 | 3 | 3 |  |
| Insertion length (bp) | 686 | 686 | 684 | 686 | 686 |  |
| Deletion count (>1 bp) | 3 | 3 | 3 | 3 | 3 |  |
| Deletion length (bp) | 4,006 | 4,006 | 4,006 | 4,006 | 4,006 |  |

\*: The strain information and original submission dates were based on best estimates from the relevant publications;
\*\*: Variant-calling was performed using CC-5325 *mt*<sup>-</sup> config 1 as the query, and previously published mitogenomic sequences as references.

Interestingly, the primary *de novo* assembly did not produce a contig corresponding to the ∼15.8-kb canonical mitogenome. To investigate this structural difference, previously published mitogenomes of *C. reinhardtii* were compiled and compared with our CC-5325 assembly (Table 1). Although the 19-kb mitogenome is approximately 21% longer than the historical reference, its overall GC content remains comparable, indicating that the expansion does not alter the nucleotide composition of the mitogenome. Across the core mitogenome, excluding the large inverted repeats, sequence variation was low, with only 10–20 SNPs and a small number of short INDELs (1–5 bp; 1–10 events) typically detected among the analyzed mitogenomes. Considering the limitations of earlier short-read sequencing technologies (9, 10), such limited sequence variation indicates that the core sequence of *Chlamydomonas* mitogenome is highly conserved. This strong sequence conservation prompted us to investigate whether the newly identified 19-kb configuration represents a genuine mtDNA structure rather than an assembly artifact.

To resolve this structural ambiguity, we examined long-read mapping profiles across the newly assembled mitogenome. Read coverage dropped sharply from 6,362× to 3,546× at nucleotide position 15,047 (Fig. 1*B*). Notably, the sequence upstream of this 15,047-bp boundary was consistent with the canonical mitogenome reference (NC_001638.1). However, the sequence extending beyond this boundary could not simply be trimmed to fit the long-standing 15.8-kb model, because individual long reads continuously spanned the full 19-kb sequence without hard or soft clipping, SNPs, or INDELs, while sustaining a robust 3,546× coverage through the terminal repeats. The single molecule sequence data demonstrated two distinct mitogenomes in CC-5325. Because the assembler we used, Flye, relies on read coverage to resolve assembly graphs (25), the overlapping sequences yet distinct read depths of the two configurations prevented simultaneous assembly of both isoforms. To recover the second configuration, we reduced the representation of HiFi reads that mapped to the 19-kb-specific regions, thereby increasing the representation of raw reads supporting the ∼15.8-kb configuration. *De novo* assembly of these filtered reads successfully generated a 15,107-bp contig. This secondary contig exhibited high, uniform read coverage and contained all expected mitochondrial gene features in a single copy, consistent with the canonical ∼15.8-kb mitogenome architecture. We designated the 19,039-bp architecture as Configuration 1 (config 1) and the 15,107-bp architecture as Configuration 2 (config 2) (Fig. 1*A*, Datasets S1−4).

These findings led us to investigate whether the dual configurations are unique to the CC-5325 strain or are broadly distributed across *C. reinhardtii* species, particularly among strains within CC-5325 lineage. This lineage traced back to the commonly used 137c mating-type minus (*mt^−^*) strain established during the 1940s–1950s, a derivative of which was subsequently deposited at the Chlamydomonas Resource Center as CC-124 (*mt^−^*). From the 137c background, the Rochaix lab isolated and selected the sister strains 4A+ (subsequently registered as CC-4051) and 17D− (unregistered) (26). In 2000, the Niyogi lab crossed 4A+ and 17D− and subsequently performed serial backcrosses to the parental 4A+, generating the strain 4A×5.2− (27). Because 4A×5.2− was estimated to share over 98% of its genetic background with the original 4A+, it was renamed as 4A− (registered as CC-4603) (26). During development of the Chlamydomonas Library Project (CLiP), the Jonikas lab crossed 4A− (CC-4603, *mt^−^*) with D66+ (CC-4425, a cell-wall-deficient *mt^+^* strain) (28). Progeny from this cross were subsequently selected to generate CC-5325, which combines genetic features derived from both parental strains (Fig. 1*C*).

To explore the distribution of the two mitogenome configurations across the CC-5325 lineage, we collected all available raw sequencing reads from related strains (17D− was unavailable) and mapped them against CC-5325 mitogenome config 1. Because the mtDNA is predominantly inherited from the *mt^−^* parent under standard crosses in *Chlamydomonas* (29, 30), we expected the CC-5325 mitogenome to show greater structural similarity to that of its *mt^−^* parent, CC-4603, than that of its *mt^+^* parent, CC-4425. However, because these datasets were generating using the 100 bp pair-end Illumina sequencing, mapping these reads to the large terminal repeats of config 1 resulted in multi-mapping. In contrast to the continuous, uniquely resolved coverage provided by our long HiFi reads, these multimapping short reads exhibited low mapping quality (MAPQ = 0) across the repeat regions (Fig. 1*B*). To validate the presence of config 1 in these strains, we focused on the critical coverage transition near the 15-kb boundary and examined read depth across the config 1-specific extended region that is absent from config 2.

Extensive reads mapping first confirmed the presence of the extended architecture in the direct *mt^−^* parent. CC-4603 showed high read coverage (4,800–7,000×) extending beyond position 15,047, providing strong support for the config 1-specific region. We then expanded this analysis to the *mt^+^* strains within the lineage. CC-4425 and CC-4051 also exhibited clear read support for this extension (50–100× and 40–600×, respectively), indicating that config 1 is not restricted to CC-5325 or its direct *mt^−^* parent, and that both config 1 and config 2 are present throughout the entire CC-5325 lineage. The substantial differences in coverage across strains further suggested variation in the relative representation of config 1. Interestingly, when mapping the raw paired-end sequencing reads from the ancestral CC-124 (*mt^−^*) and CC-125 (*mt^+^*) strains to config 1, we also identified substantial read coverage across the extended region (400–2,000× for CC-124 and 1,200–2,500× for CC-125) (*SI Appendix*, Fig. S1). Because CC-124 and CC-125 represent two early laboratory lines of *C. reinhardtii*, these results suggest that the presence of two mitogenome configurations predates the derivation of CC-5325 and may be broadly distributed among laboratory strains.

### The canonical structure of the mitogenome termini is not supported in CC-5325

Previous models proposed that the linear mitogenome of *C. reinhardtii* possesses terminal telomeric structures characterized by 532-bp inverted repeats (10). However, mapping our long reads to the reference mitogenome (NC_001638.1) revealed substantial structural discrepancies at its termini. While the core genomic region exhibited high and continuous mapping coverage with high sequence fidelity, the terminal regions lacked continuous long-reads support and displayed abrupt coverage spikes (*SI Appendix*, Fig. S4*A*). Sequence analysis also revealed no tandem repeats that were similar to the chromosomal telomeric arrays of *Chlamydomonas* (5′-TTTTAGGG-3′). Reanalysis of the historical inverse PCR data (GenBank: M83998.1) further indicated that the previously reported terminal fusions could reflect either a template-switching PCR artifacts mediated by a 12-bp homologous motif or internal junctions derived from a larger concatemeric mtDNA molecule (detailed in *SI Appendix, Supporting Information Text*).

### Structural resolution reveals a highly dynamic, multipartite mitogenome architecture

To further explore the structural heterogeneity of the mitogenome, we extracted all raw HiFi and ONT reads longer than 19 kb that mapped to the config 1 and config 2 architectures and examined their native conformations. This single-molecule profiling revealed a complex population of discrete monomers and multimeric concatemers in different arrangements of the two configurations. Full-length monomeric reads were recovered that spanned and closely matched both configurations 1 and 2, providing direct read-level support for the assembled structures (Fig. 2*A*, upper panel). Config 1 was supported by ONT and HiFi reads at sequencing depths of 2,588× and 155×, respectively; config 2 similarly exhibited strong read support, with ONT and HiFi coverage of 378× and 259×, respectively. The longer ONT mitogenome reads (N50 = 23,508 bp), compared with the HiFi reads (N50 = 21,735 bp), enabled direct resolution of a diverse array of multimeric concatemers. The most abundant, high-confidence concatemeric structures were [config 2–config 2] and [config 1–config 2] concatemers. Furthermore, complex higher-order arrangements with three or four interconnected configurations were discovered, revealing an extensive diversity of concatemeric mtDNA structures (Fig. 2*A*, lower panel). Together, the recovery of monomeric and multimeric mtDNA molecules provides a snapshot of substantial structural heterogeneity and is consistent with ongoing transitions among alternative mtDNA configurations, which may represent recombination intermediates associated with mtDNA replication. Notably, the fusion PCR product previously reported by Ma et al. (31) may represent a junction between these concatemers, such as those observed in [config 1–config 2] or [config 2–config 2] fusions (*SI Appendix*, Fig. S4*C*).

**Fig. 2.**
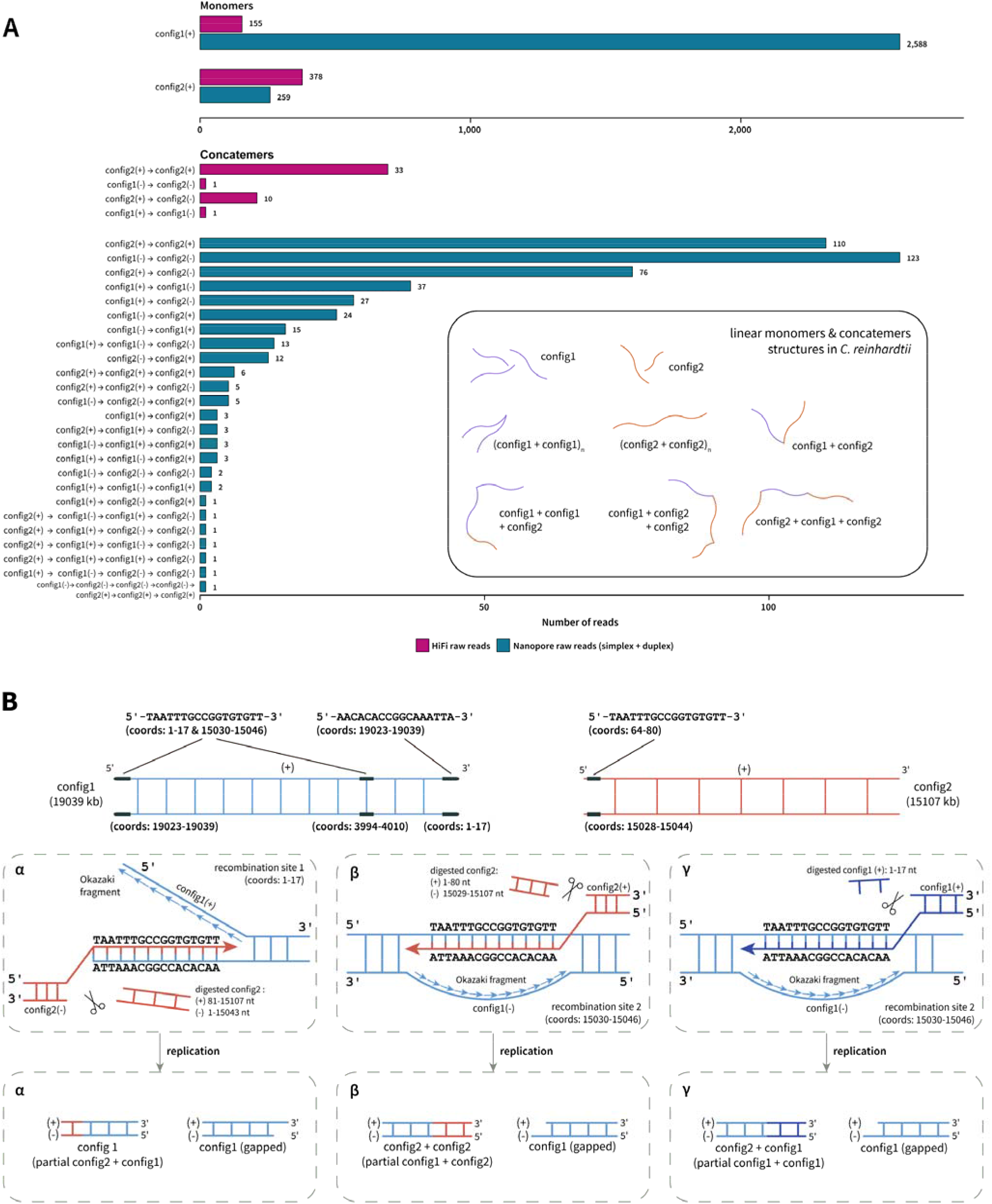
Long-read sequencing reveals mitogenome heteroplasmy and structural dynamics in *C. reinhardtii*. (A) Quantification and structural classification of mitogenome monomers and concatemers identified using PacBio HiFi (magenta) and Oxford Nanopore Technologies (ONT; teal) long-read sequencing. Bar charts detail the specific read counts supporting various structural arrangements involving config 1 and config 2. The inset box schematically illustrates the diverse linear monomeric and concatemeric mtDNA structures detected by long-read sequencing. (B) Proposed recombination-dependent replication (RDR) model driving mitogenome structural diversity. The schematic outlines three proposed recombination and replication pathways (modes α, β, and γ) involving a shared 17-bp homologous sequence in config 1 and config 2 that could contribute to the generation of diverse mitogenome configurations.

### Sequence homology of a conserved 17-bp motif supports a recombination-dependent replication (RDR) model

ONT long reads revealed that the multimeric concatemers of the CC-5325 mitogenome were not strictly organized as simple, head-to-tail tandem fusions of the config 1 and config 2. Instead, most of these inter-genomic reads shared a highly conserved 17-bp consensus sequence motif (“seam“: TAATTTGCCGGTGTGTT, reverse complement AACACACCGGCAAATTA) at the junctions between configurations (Dataset S5). Mapping this seam across both configurations revealed its precise genomic coordinates: on the sense (+) strand of config 1, the forward 17-bp motif occurred at the left terminal region (coordinates 1–17) and internally (coordinates 15,030–15,046), while its reverse complement occurred at the right terminal region (coordinates 19,023–19,039). On the sense (+) strand of config 2, a single copy of forward 17-bp motif occurred near the 5′ end at coordinates 64–80 (Fig. 2*B*).

Because linear organellar genomes face the end-replication problem, we propose that the spatially separated homologous 17-bp motifs enable config 1 to serve as a replication template. By providing homologous sites for strand invasion or so called “landing pads,” this architecture could facilitate a continuous Recombination-Dependent Replication (RDR) mechanism. In the primary invasion mechanism (Fig. 2*B*, mode α), we propose that replication could be initiated by nucleolytic resection (via exo- or endonucleases) of the double helix to expose the 17-bp homology motif as a single-stranded 3′-OH “sticky end.” This 3′ end could then invade the complementary sequence in the negative strand of config 1, providing a primer for DNA polymerase-mediated strand-displacement synthesis towards the terminus. Resolution of the resulting Holliday junction (32) could separate the intermediate into distinct molecules: the uninterrupted leading strand seamlessly forms the concatemer, while the lagging strand forms a new template molecule. Importantly, the end-replication problem on the newly synthesized lagging strand could leave a 5′ gap. This incomplete synthesis naturally may expose a single-stranded 3′-OH overhang on the complementary negative strand. Rather than representing a terminal replication product, this 3′ overhang could provide a substrate to prime subsequent strand invasion, driving iterative cycles of synthesis and junction resolution that generate the observed multimeric mtDNA structures.

These invasion mechanisms may be highly versatile and operate across multiple genomic orientations (Fig. 2*B*, modes β and γ). For instance, in the mode β, exonucleolytic restriction of the 5′ terminus (coordinates 1–80) of the config 2 (+) strand could expose the complementary (-) strand 17-bp motif. Concurrent nucleolytic removal of the nonhomologous terminal sequence (coordinates 15,029–15,107) could eliminate overhanging flaps that would otherwise cause steric hindrance during subsequent elongation, thereby generating a compatible substrate for strand invasion at the homologous site in the config 1 (+) template (Fig. 2*B*, modes β). Following polymerase extension and Holliday junction resolution, this proposed pathway could generate a hybrid [config 2–17 bp motif–config 2] concatemer while leaving a gapped config 1 with exposed 3′-end that could participate in subsequent rounds of strand invasion. Furthermore, in mode γ, config 1 molecules could utilize a parallel RDR pathway involving strand invasion between config 1 molecules. Because the 17-bp motif is located at the extreme terminus of config 1 (coordinates 1–17), RDR initiation via this pathway would require only limited 5′-end nucleolytic resection of the first 17 nucleotides to directly expose the complementary 17-bp sequence as a single-stranded 3′ overhang with a free 3′-OH on the negative strand, thereby providing a substrate for homologous strand invasion (Fig. 2*B*, modes γ). These continuous, multidirectional homologous interactions could generate [config 2–17 bp motif–config 1] concatemers while regenerating config 1 molecules containing an exposed 3′-end capable of driving successive rounds of RDR.

Importantly, the replication models proposed in Fig. 2*B* may not occur as isolated, sequential events involving discrete monomeric molecules. Instead, multiple RDR events could occur concurrently among monomers, subgenomic fragments, and concatemers, potentially generating an interconnected and actively recombining population of mtDNA molecules (Dataset S5). While the branched and concatemeric mtDNA structures we observed are well explained by this RDR model, how these complex mtDNA intermediates are resolved and segregated, and how transcriptional machinery navigates such a hyper-branched, structurally heterogeneous DNA population, remain important questions for future investigation.

Strong support for our proposed RDR model came from a specific structural constraint imposed by the architecture of the 19 kb config 1. The geometry of the config 1 terminal repeats and the predicted directionality of the proposed RDR model preclude formation of a linear, head-to-tail fusion of a [config 1–17 bp motif–config 1] concatemer (Fig. 2*B*). Consistent with this prediction, exhaustive examination of our ONT reads (877 concatemers in total) identified no molecules with this specific structural arrangement (Dataset S5). The absence of this theoretically prohibited concatemer provides independent structural support for the directional and geometric constraints predicted by the RDR model.

Despite this structural support, our RDR model primarily accounts for config 1-driven concatemeric replication and did not explain how config 2 replicates autonomously in the absence of config 1 or the 17-bp motif. Determining whether mitogenome config 2 can be maintained as an independent architecture requires evaluating mitogenomic structural diversity across additional *Chlamydomonas* lineages.

### Prevalent homology exists between nuclear and mitogenome in the CC-5325 lineage

Based on the complete telomere-to-telomere CC-5325 nuclear genome and the newly resolved mitogenome, we reevaluated sequence homology between the nuclear and mitochondrial genomes of *C. reinhardtii*. We identified a widespread but highly unevenly distributed nuclear regions with sequence homology to the mitogenome (Fig. 3). These homologous fragments were generally short—typically ranging from 30 to over 300 bp—but frequently exceeded 90% identity between the nuclear and mitochondrial sequences (*SI Appendix*, Table S1). These sequences included both intergenic regions and portions of conserved mitochondrial genes, including *cob* and *nad5*. However, their distribution across the nuclear genome was highly heterogeneous. Mapping these homologous sequences likely nuclear mitochondrial DNA segments (NUMTs), across the nuclear chromosomes revealed particularly high representation on chromosomes 01, 02, 09, 16, and 17. By contrast, chromosome 11 contained no mitochondrial sequence match. Although sequence similarity alone does not establish the direction of DNA transfer between the nuclear and mitochondrial genomes, this spatial distribution is inherently informative and raises a question whether this pattern is conserved among related strains in the *C. reinhardtii* CC-5325 lineage.

**Fig. 3.** Sequence homology between the mitogenome and nuclear genome of *C. reinhardtii* CC-5325. The Circos plot shows the distribution of homologous sequences between mitogenome config 1 (top-left quadrant) and 17 nuclear chromosomes (Chr_01 to Chr_17; blue arcs). The outer mitogenome track displays annotated functional features, including coding genes (yellow), ribosomal RNAs (blue), transfer RNAs (pink), and pseudogenes (brown). The inner mitogenome track distinguishes the non-repeating region (black) from the terminal inverted repeats (white). Green ribbons connect specific mitochondrial loci to their homologous regions distributed across the nuclear chromosomes.

To further assess sequence homology among the different genomic compartments, we next examined mtDNA sequences homologous to the plastome, including mitochondrial plastid DNA segments (MTPTs). We identified only one tRNA with complete identity, trnQ-UUG, and limited homologous regions within two rRNAs (*rrnL7* and *rrnL8*) (Fig. 4), indicating the sequence homology between the plastome and mitogenome is relatively low in *C. reinhardtii*.

**Fig. 4.**
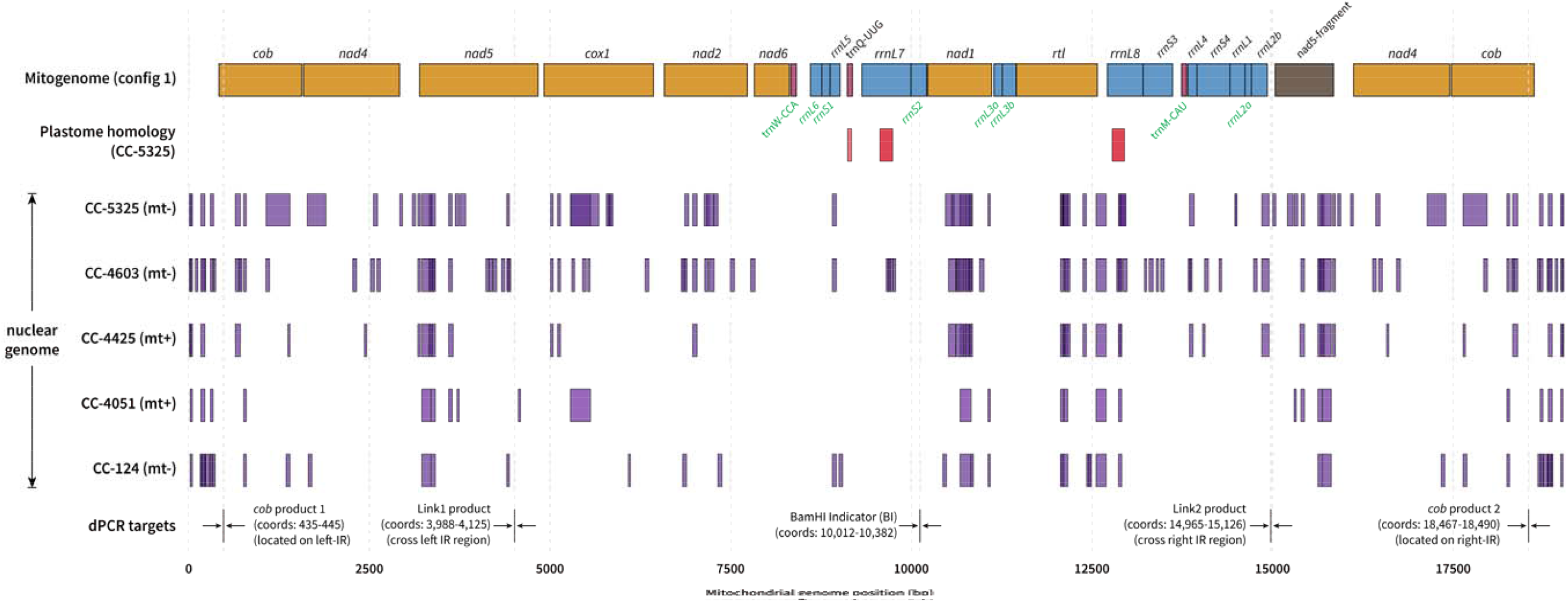
Intergenomic sequence homology and dPCR assay design across the *C. reinhardtii* CC-5325 lineage. The top track displays the annotated linear structure of the CC-5325 mitogenome config 1, which serves as the reference for genomic coordinate. The second track (red) identifies regions of sequence homology between mitogenome config 1 and the CC-5325 plastome. The five subsequent tracks (purple) show the distribution of intergenomic sequence homology between mitogenome config 1 and the nuclear genomes of five related strains [CC-5325 (*mt^−^*), CC-4603 (*mt^−^*), CC-4425 (*mt^+^*), CC-4051 (*mt^+^*), and CC-124 (*mt^−^*)]. The bottom track indicates the genomic coordinates of four dPCR amplicons [*cob*, link1, BamHI Indicator (BI), and Link2] (*SI Appendix*, Table S4). These amplicons were designed to avoid regions with nuclear and plastid sequence homology, minimizing potential off-target amplification. Link1 amplifies a region shared by config 1 and config 2, whereas Link2 targets a boundary region that is config 1-specific.

Prompted by the non-random distribution of NUMTs in CC-5325, we next investigated whether this genomic pattern was conserved across the CC-5325 lineage or was a transient, strain-specific feature to CC-5325. Analysis of related *mt^−^* strains, including CC-4603 and CC-124, revealed that the distribution pattern of NUMTs was highly conserved. These strains exhibited widespread mitochondrial sequence homology within the nuclear genome, encompassing both mitochondrial intergenic regions and most mitochondrial core genes. However, a specific subset of genome features—including *nad6*, trnW-CCA, trnM-CAU, and several rRNA subunits (*rrnL6*, *rrnS1*, *rrnS2*, *rrnL3a*, *rrnL3b*, *rrnS3*, *rrnS4*, *rrnL2a*)—showed no nuclear matches (Fig. 4). Although we observed lower NUMT representation in CC-124, this discrepancy unlikely reflected a biological loss during the subsequent laboratory derivation of the CC-5325 lineage (Fig. 1*C*). Instead, it likely reflected differences in genome assembly completeness: the raw data for CC-124 genome was assembled from short-read Illumina data and covers only ∼75% of the T2T-level CC-5325 genome (*SI Appendix*, Table S2). Despite this incomplete assembly, the ancestral CC-124 strain retained overall distribution pattern of mitochondrial–nuclear sequence homology, indicating that these shared sequences were already present in their common ancestral background and predated the subsequent derivation of the CC-5325 lineage.

Most notably, this widespread NUMT pattern was not restricted to the *mt^−^*lineage, despite the predominant inheritance of mitochondrial DNA from the *mt^−^*parent in *Chlamydomonas*. The *mt^+^* strains CC-4425 and CC-4501, exhibited a strikingly similar NUMTs distribution pattern across coding, ribosomal, and intergenic regions compared to those of the *mt^−^* strains. Although a few NUMT fragments were absent in these *mt^+^* assemblies—potentially reflecting incomplete sequencing coverage (*SI Appendix*, Table S2) or sequence divergence among strains—the overall NUMT pattern remained largely conserved. Together with the conservation observed in CC-124, these results indicate that this NUMT distribution is broadly shared among *mt^−^* and *mt^+^* strains and has persisted through the laboratory crosses that generated the CC-5325 lineage (Fig. 1*C*).

### Mitogenome configuration abundance varies across the CC-5325 lineage

Given the coexistence of two distinct mitogenome configurations, we next determined their relative abundance across different genetic backgrounds. To quantify the relative abundance of each configuration, we utilized a complementary approach combining sequence-depth analysis with experimental validation by digital PCR (dPCR).

Analysis of whole genome sequencing (WGS) read mapping profiles revealed that config 1 to config 2 were present at comparable abundances in CC-5325 with config 1 representing an estimated fraction of 0.56 (config 2 fraction of 0.44, Fig. 5*L*). Utilizing high-quality long-read sequence data spanning the 15,047-bp boundary of config 1 (Fig. 1*B*), we designed dPCR assays targeting the Link1 (representing the total mitogenome pool) and Link2 (config 1-specific) regions, with a BamHI-indicator (BI) amplicon serving as an internal reference (Fig. 4). In CC-5325, the Link1 copy number was comparable to that of the BI reference, and the calculated relative abundance of config 1 (Link2/Link1) was 0.54 ± 0.04 (Fig. 5*A*). This close agreement between the dPCR measurement and the *in silico* read-coverage estimate independently supported the accuracy of both approaches and further support the structural model of configs 1 and 2.

**Fig. 5.**
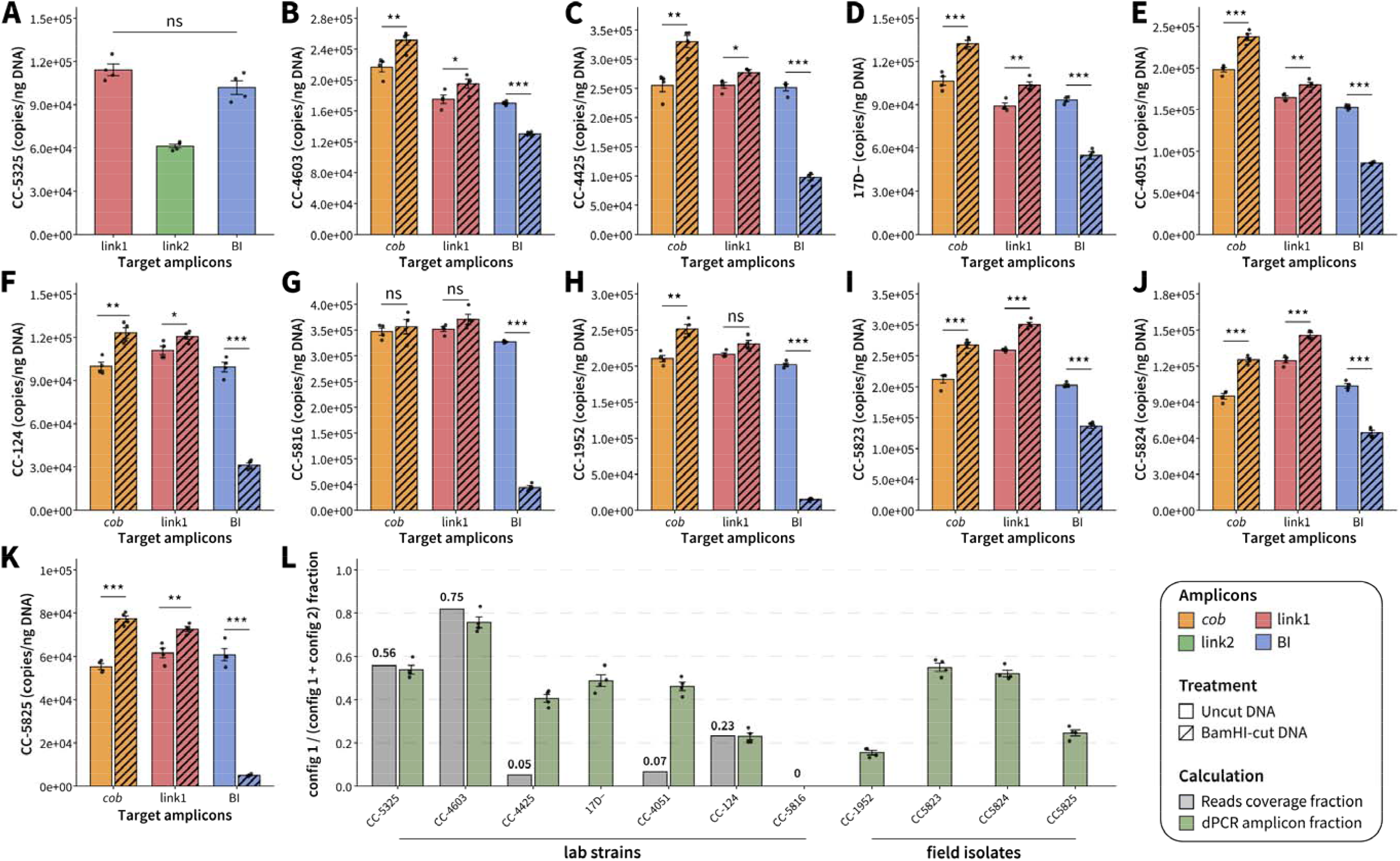
Absolute quantification of mitogenome configurations using dPCR. Copy number of four target amplicons (Link1, Link2, BamHI indicator [BI], and *cob*) were used to quantify mtDNA and distinguish mitogenome configurations. (A) Direct amplicon quantification in strain CC-5325, using Link1, Link2, and BI to estimate the abundance of total mitogenome (config 1 + 2), config 1, and the total mitogenome pool (internal control), respectively. (B–K) dPCR quantification in related CC-5325 lineage strains (B-F), a negative control (G), and field-isolated strains (H-K). For these strains, configuration abundances were estimated from changes in detectable target concentrations following BamHI digestion. The increase in *cob* signal (Δ*cob*–Δlink1) reflects the config 1 monomer-specific contribution, whereas the decreases in BI signal (ΔBI–Δlink1) represents the cleavage of BI-containing sequences associated with the total pool of config 1 and config 2. More details about dPCR quantification strategies are included in *SI Appendix, Fig. S8*. (L) Comparison of config 1 and config 2 relative abundance estimated from *in silico* genomic read coverage and *in vitro* dPCR measurements. Statistical significance: *, *P* < 0.05; **, *P* < 0.01; ***, *P* < 0.001; ns, not significant.

In contrast to CC-5325, short-read WGS mapping from its *mt^−^* parent, CC-4603—the expected source of the CC-5325 mitogenome through uniparental inheritance—yielded a high config 1 fraction of 0.75 (Fig. 5*L*). Conversely, the *mt^+^* strains in the lineage displayed substantially lower estimated config 1 fractions (0.05 in CC-4425 and 0.06 in CC-4051), and the ancestral *mt^−^* strain CC-124 showed a fraction of 0.23 (Fig. 5*L*). We found these short-read-based estimates were potentially confounded by INDELs and larger structural variants concentrated around the 15,047-bp boundary in all strains except CC-5325 (Fig. 1*B*). This localized sequence divergence precluded the design of reliable Link2 amplicon primers for the non-CC-5235 strains. To circumvent this limitation, we utilized the BI amplicon (Fig. 4) as a reference for the total mitogenome pool prior to restriction digestion. Following BamHI digestion and dPCR, we observed a significant increase in the copy numbers of both *cob* and Link1 (Fig. 5*B*–*F* and *H*–*K*). This digestion-induced increase in Link1 copy number provides independent experimental support for the presence of multimeric concatemers in CC-4603, CC-4425, 17D–, CC-4051, and CC-124: if configs 1 and 2 existed exclusively as independent monomers, BamHI digestion would not affect the measured Link1 copy number; conversely, BamHI cleavage of concatemers can separate physically linked multimeric mtDNA molecules, allowing them to partition independently during dPCR and thereby increasing their measured copy number even when digestion is incomplete (*SI Appendix*, Fig. S8). These findings corroborate our observations in CC-5325 (Fig. 2*A*) and further demonstrate that complex concatemeric mitogenome architectures are broadly distributed across the CC-5325 lineage.

Next, we applied this restriction-coupled, multi-target differential dPCR strategy (integrating Δ *cob*, Δ Link1, and Δ BI; see Materials and Methods) to quantify the relative abundance of config 1 to config 2 ratios across the broader lineage. Our experimental data revealed that CC-4603 had a config 1 fraction of 0.76 ± 0.05, representing the highest proportion among the strains analyzed (Fig. 5*L*). Tracing the mitogenomic inheritance lineage, the intermediate *mt^−^* parent 17D− exhibited a relatively balanced config 1 fraction (0.49 ± 0.05; Fig. 5*D*), whereas the ancestor strain CC-124 displayed a comparatively low config 1 fraction (0.23 ± 0.03; Fig. 5*F*). Substantial variation was also observed among the *mt^+^* strains, with CC-4425 and CC-4051 showing config 1 fractions of 0.41 ± 0.04 and 0.46 ± 0.04, respectively (Fig. 5*C, E*). The config 1 fraction showed no simple relationship with strain history or mating type, which could reflect stochastic segregation or effects of the extensive laboratory crossing and backcrossing within this lineage. Crucially, this differential dPCR measurement corroborated *in silico* mapping estimates for CC-4603 and CC-124, in which structural variation near the 15,047-bp boundary is minimal. Moreover, dPCR measurements successfully corrected the config 1 fractions for CC-4425 and CC-4051, in which localized sequence divergence had severely underestimated config 1 fractions using *in silico* short-read mapping (Fig. 1*B*, Fig. 5*L*).

### Mitogenome structural heteroplasmy varies across diverse *C. reinhardtii* laboratory lineages and field isolates

To assess the broader prevalence of mitogenome structural heteroplasmy, we analyzed three strains with available long-read sequencing data (CC-4532, CC-5816, and UL-1690) that lie outside the CC-5325 lineage but share ancestry with CC-124 (Levene’s 137c line) (*SI Appendix*, Fig. S2). Mapping these reads to config 1 revealed reads in CC-4532 that spanned the diagnostic 15,047-bp boundary. Although coverage across this region was relatively low (60–150×), the presence of junction-spanning reads supports the occurrence of config 1 in CC-4532. Conversely, no reads spanning the diagnostic config 1 boundary were detected in CC-5816 and UL-1690 (*SI Appendix*, Fig. S3), indicating config 1 may not be present in these strains. *De novo* assembly of the CC-5816 mitogenome yielded a single predominant configuration with an IR (468 bp) – Core Region (14,722 bp) – IR (468 bp) architecture (*SI Appendix*, Fig. S7), supported by near-uniform read coverage and the absence of substantial soft-or hard-clipped reads (*SI Appendix*, Fig. S5*B*). This discrete 15,664-bp assembly (Dataset S6) structurally resembles the architecture of CC-5325 config 2 and the canonical mitogenome (*SI Appendix*, Fig. S5*A*). Because *in silico* read mapping detected no evidence of config 1, CC-5816 served as a negative control for our dPCR quantification system. Consistent with this result, *cob* amplicon levels did not significantly increase following BamHI digestion, providing independent experimental support for the absence of config 1 in CC-5816 (Fig. 5*G*).

Interestingly, our proposed RDR model for CC-5325 provides no obvious pathway for autonomous replication of config 2 (Fig. 2*B*). However, the predominance of config 2 monomers in CC-5816 necessitates a mechanism for its autonomous replication independently of detectable config 1. Structurally, the CC-5816 mitogenome contains 468-bp terminal inverted repeats, which could provide homologous sequence substrates or “landing pad” for RDR model (*SI Appendix*, Fig. S7). Despite this structural compatibility, systematic analysis of CC-5816 ONT reads (Dataset S7) failed to identify a recurrent homologous sequence motif comparable to the highly conserved “17-bp seam” proposed to drive RDR in CC-5325 (Dataset S5). Therefore, whether its replication involves an RDR-like mechanism remains unresolved. Further investigation will be required to identify recombination sites and determine the mechanism underlying replication of monomeric config 2 in CC-5816.

To determine whether the observed mitogenome structural heteroplasmy extends beyond laboratory-derived lineages, we evaluated four field-isolated strains collected from distinct geographical regions and years (CC-1952, CC-5823, CC-5824, and CC-5825) (*SI Appendix*, Table S5). Quantification of configuration proportions revealed substantial variation among these field isolates. CC-1952 exhibited the lowest config 1 fraction recorded across all analyzed strains (0.16 ± 0.02), whereas CC-5825 showed a fraction (0.25 ± 0.03) comparable to that of the laboratory strain CC-124 (Fig. 5*L*). By contrast, CC-5823 and CC-5824 displayed nearly balanced configuration proportions (0.55 ± 0.04 and 0.52 ± 0.03, respectively) (Fig. 5*L*). Thus, substantial variation in the relative abundance of config 1 and config 2 is not restricted to the laboratory-derived lineage but also occurs among geographically and temporally distinct field isolates. Because these field isolates were exempt from laboratory crossing and subsequent selection process, these observations suggest that mitogenome heteroplasmy and variable configuration abundance across *C. reinhardtii* strains is not readily explained by laboratory crossing history alone.

## Discussion

### Overcoming algorithmic constraints to resolve mitogenome architecture

Organelle genome assembly has often been considered as a relatively straightforward task, given the small size, relatively low repeat contents, and highly conserved gene content of mitochondrial and plastid genome (33). Conserved organelle-encoded genes such as *rbcL* and *COI* are wildly used as taxonomic markers (34); however, sequence conservation does not necessarily imply conservation of organelle genome architecture. Driven by the prevalence of short-read sequencing technologies, conventional organelle genome assembly pipelines have generated primary unitigs that are often iteratively extended until they align with a reference template or circularize (35). Consequently, organelle assemblies relied on scaffolding against related model species may failed to resolve structurally divergent contigs or exclude them as chimeric artifacts. Such limitations can bias the resulting assemblies toward established reference architectures, potentially obscuring genuine structural variation and reducing heterogeneous (mito)genomic DNA populations to an idealized single consensus representation.

Recently developed long-read assemblers, such as Flye (25) and Hifiasm (36), have substantially improved the resolution of complex genomic repeats. However, they operate under strict ploidy and read-depth assumptions designed for nuclear chromosomes (37). These tools resolve assembly graphs by calculating average coverage to compute a single contiguous path (38). Because multiple structural conformations may co-exist *in vivo*, resolving these branching graphs into a single path can obscure alternative configurations and their relative coverage (or abundance). Our long-read analysis of the *C. reinhardtii* CC-5325 mitogenome directly illustrates this limitation. While the overall gene content remains conserved relative to earlier reference sequences, we uncovered substantial differences in mitogenome architecture (Fig. 1*A*). Maintaining identical coding sequences across discrete, alternative architectures presents a key challenge for conventional assembly algorithms. Because these pipelines prioritize resolving overlapping reads into a contiguous path (contig) (39), alternative structural configurations are routinely merged into one. In our initial *de novo* assembly using default parameters, only config 1 was recovered whereas the alternative config 2 was not. We subsequently recovered config 2 by restricting the coverage parameters, preventing the algorithm from collapsing the data into config 1. Recognizing these limitations, we bypassed the assembly process with direct analysis of individual raw long reads, often longer than a single genome to characterize alternative mtDNA configurations and concatemeric structures.

Because ONT sequencing analyzes long, unamplified DNA molecules directly, the resulting reads can preserve their long-range sequence feature and provide direct single-molecule evidence of mtDNA organization (40). By contrast, PacBio HiFi sequencing generates highly accurate consensus sequences through multiple passes of circularized DNA templates prepared by DNA fragmentation and adapter ligation (41). While this approach yields highly accurate sequence data ideal for resolving mitochondrial gene content and structural configurations, the DNA fragmentation step does not preserve the native structural topology of the mtDNA molecules *in vivo*. Direct mapping of raw ONT reads revealed both linear monomeric and concatemeric mtDNA structures in *C. reinhardtii*, providing evidence of substantial structural heterogeneity within the mtDNA population (Fig. 2*A*), which aligns strongly with early electron microscopy studies reporting extended and supercoiled mtDNA architectures (3). Nevertheless, although ONT sequencing provides single-molecule information on mtDNA architectures, it represents only a snapshot of the DNA molecules recovered at the time of sampling, but cannot capture temporal changes in mtDNA architecture. Although our results revealed substantial variation in the relative abundance of configs 1 and 2 across different *Chlamydomonas* strains with distinct genetic backgrounds (the CC-5325 lineage and related strains, as well as four field strains outside this lineage), whether these proportions also vary in response to cellular physiological conditions remains unknown. To control for this uncertainty, all strains analyzed by dPCR were all sampled during stationary growth phase, whereas growth phases and culture conditions associated with the publicly available sequencing datasets retrieved from the NCBI Sequence Read Archive (SRA) (Fig. 1*B*) remain unclear.

### Mitogenome structure heteroplasmy beyond transient biparental inheritance

Although transient mitochondrial heteroplasmy occurs following zygote formation in *C. reinhardtii*, persistent heteroplasmy has generally been considered uncommon because mtDNA inherited from the *mt^+^* parent is rapidly eliminated, typically within 3–6 h after fertilization (42). In this study, for the first time, we provide evidence of persistent structural heteroplasmy in *C. reinhardtii* by defining distinct mitogenome configurations and concatemeric arrangements, quantifying their copy numbers, and documenting their coexistence across the CC-5325 lineage and field-isolated strains (Figs. 2 and 5).

In angiosperms, mitochondrial DNA is predominantly maternally inherited, however, persistent heteroplasmy driven by *de novo* mutations or occasional paternal leakage has been documented (16). For example, in *Silene vulgaris*, paternal leakage has been shown to produce heteroplasmic offspring harboring both maternal and paternal mitochondrial genotypes (18). Furthermore, subsequent crossing studies demonstrated that maternal transmitted heteroplasmy could persist across generations, although the evenness of mitochondrial variant representation usually declined, with heteroplasmy frequently lost in subsequent generations through vegetative segregation (43). These observations demonstrate that mitochondrial heteroplasmy is frequently observed in land plants, although the relative abundance of mitochondrial variants may fluctuate and heteroplasmy can eventually be lost in their offspring. A comparable pattern of variation in mitochondrial genome composition is evident in *C. reinhardtii*, where our results reveal the persistence of alternative mitogenome configurations in the CC-5325 lineage (Fig. 5), together with substantial variation in their relative abundance across related strains (*SI Appendix*, Fig. S3). In land plants, vegetative segregation can alter mitochondrial variant frequencies through stochastic partitioning, a process analogous to genetic drift that may ultimately result in cellular homoplasmy (44). However, the initial dynamics of heteroplasmy differ fundamentally between these systems. In most angiosperms, paternal mtDNA is confined to the pollen and physically excluded prior to or during fertilization, preventing parental mitogenomes from ever mixing (16). Conversely, *C. reinhardtii* fertilization involves the complete fusion of *mt^+^* and *mt^−^* cells, forcing a transient but direct intracellular coexistence of both parental mitogenomes. Given this unique zygotic environment, whether the persistence or loss of mitogenome config 1 in *C. reinhardtii* is governed primarily by stochastic vegetative segregation, differential replication, or other molecular mechanisms remains an important unresolved question.

While targeted genetic markers (e.g., SNPs) have been used to document persistent heteroplasmy in land plants (16), the specific mechanisms underlying paternal leakage and mtDNA segregation remain difficult to resolve using these markers alone. Recent advances in the yeast *Saccharomyces cerevisiae* have combined mtDNA-based fluorescent markers, microfluidics, and automated single-cell tracking to investigate mitochondrial inheritance dynamics *in vivo* (24). In wild-type yeast cells, mitochondrial fission and fusion contribute to the redistribution of mtDNA variants within the mitochondrial network. During budding, daughter cells inherit a subset of the parental mitochondrial population, leading to heteroplasmy for an extended period (45). By contrast, when mitochondrial fission is disrupted (e.g., through *DNM1* deletion), mtDNA shuffling is impaired, potentially accelerating the segregation of distinct mtDNA variants. In the absence of normal mitochondrial fission, newly replicated mtDNA molecules might remain spatially associated with their parental templates rather than being efficiently redistributed throughout the mitochondrial network. This restricted spatial redistribution could promote the inheritance of clusters enriched in particular mtDNA variants, thereby increasing segregation bias and accelerating the transition toward homoplasmy (24). These mechanistic insights from yeast provide a framework for understanding the persistence and segregation of the two distinct mitogenome configurations identified in *C. reinhardtii*. It is possible that mitochondrial dynamics and associated regulatory proteins may similarly influence the maintenance, abundance, and segregation of config 1 and config 2 in *C. reinhardtii*, potentially during zygote formation and maturation processes, a hypothesis that warrants further investigation.

### Alternative mitogenome configurations suggest distinct replication mechanisms

Historically, Vahrenholz et al. proposed an *rtl*-dependent replication model for the linear mitochondrial genome of *C. reinhardtii*, hypothesizing that its terminal regions contain 3′ single-stranded overhangs that could be primed by RNA derived from an internal telomeric repeat and subsequently elongated by a reverse transcriptase-like (RTL) enzyme (10). To address the end-replication problem associate with linear genomes, this model proposed that the RTL enzyme complements and extends mitochondrial DNA termini. A recent comparative genomic study demonstrated that *rtl* gene was present but may not be universally required for mitogenome replication among closely related green algae, particularly in species with circular-mapping mitogenomes or terminal hairpin architectures (11). Furthermore, the classical *rtl*-based model invokes an 86-bp region of sequence homology between both termini and an internal mitogenomic locus. In this study, we found that the terminal sequences of the *C. reinhardtii* CC-5325 config 2 and CC-5816 mitogenomes differ from those described in the classical model and lack homology to the internal mitogenomic sequences (*SI Appendix*, Fig. S5*A*). The absence of the proposed terminal-internal sequence homology may limit the applicability of the classical *rtl*-mediated replication model, which invokes an RNA primer derived from the homologous internal sequence. We further confirmed that the assembled terminal regions were validated by sequence reads without evidence of soft-clipping artifacts (*SI Appendix*, Fig. S5*B*), supporting variation in mitogenome terminal architecture among *C. reinhardtii* strains. Thus, the classical *rtl*-based model may not fully explain mitochondrial genome replication across different *C. reinhardtii* lineages, particularly those harboring config 2-like mitogenomes and lacking the terminal-internal homology proposed in the original model. Consequently, beyond structural heteroplasmy, *C. reinhardtii* mitogenomes may exhibit heterogeneity in mitochondrial DNA replication mechanisms driven by the presence of config 1. In lineages harboring config 1, the observed concatemeric arrangements are consistent with the proposed RDR model; conversely, in lineages lacking config 1, their mitogenome topology precludes RDR, alternative replication or terminal maintenance mechanisms may contribute to the persistence of config 2-like mitogenomes.

Notably, early attempts by Vahrenholz et al. to clone mitochondrial terminal fragments encountered difficulties with conventional plasmid ligation—a limitation driven by the requirement for blunt or highly specific cohesive termini of the vector (10, 46). These historical observations are also compatible with our proposed RDR model. Rather than necessarily representing static structural features of mature linear mitogenomes involved in *rtl*-mediated extension, the reported exposed 3′ single-stranded overhangs could represent transient recombination intermediates of config 1 (*Fig. 2B*) within a population of heterogeneous, replicated and replicating mitochondrial DNA. Although our RDR model provides a possible explanation for the occurrence of these single-stranded overhangs, the predicted 17-nt sequence at the 3′ extension differs from the 41-nt overhang reported previously (10).

This discrepancy raises an important mechanistic question for our modified RDR model: How does *C. reinhardtii* enzymatically process, digest, or resolve these replicating monomers and concatemeric junctions *in vivo*? Previously proposed models, involving T-loop formation or DNA folding may also require site-specific endonuclease activity (11). However, no gene encoding an enzyme with the proposed junction-processing activity has been found in the *C. reinhardtii* mitogenome. Addressing this question is challenging because previous studies of organellar endonucleolytic activity in chlorophytes has focused on post-transcriptional RNA maturation (47). Nevertheless, recent studies have shown that experimentally induced reduction of mitochondrial DNA copy number triggers retrograde signaling associated with increased expression of nuclear-encoded respiratory genes including *AOX1* (48). Given the crucial role of the nuclear-encoded surveillance machinery in balancing plant mitochondrial DNA replication, recombination, and repair (49), enzymes involved in processing RDR intermediates may be dynamically regulated in response to retrograde signals tracking mitogenome abundance, structural integrity, or stoichiometric shifts. Together, the mechanisms of mtDNA-end processing and concatemer resolution, the identities of the responsible enzymes, and their potential regulation by mitochondrial signaling in our modified RDR model remain unresolved and warrant targeted genetic and biochemical investigation.

### Underestimated prevalence and dynamic distribution of NUMTs in *C. reinhardtii*

Because isolating highly purified mitochondrial DNA from *C. reinhardtii* is technically challenging (3), many mitochondrial gene and transcriptomic analyses rely on total cellular nucleic acids as templates. This approach introduces the risk of inadvertently amplifying NUMTs rather than authentic mtDNA sequences. Previous comparative analyses indicated that organellar-derived DNA insertions into the nuclear genome of *C. reinhardtii* were rare, in contrast to the extensive arrays of NUMTs documented in higher plants and mammals (50). However, these earlier assessments may have been constrained by incomplete or fragmented genome assemblies and the sensitivity of the sequence alignment algorithms employed. Leveraging a recently published gap-free nuclear genome assembly, a genomic analysis of CC-5816 identified 56 organellar-derived nuclear DNA insertions, including 23 distinct NUMTs exhibiting 87.8% to 100% sequence identity to mtDNA (51). This discovery further supports our conclusion that NUMTs are more prevalent in *Chlamydomonas* than previously recognized. Interestingly, CC-5816—a strain derived from a single-colony isolate following a cross between CC-125 and CC-124—exhibits no NUMT insertion on chromosomes 4, 8, 9, 11, 12, or 14. This distribution differs from the chromosomal distribution of NUMTs observed in the CC-5325 lineage (Fig. 3). This intraspecific variation in NUMT distributions among *C. reinhardtii* lineages may reflect intracellular gene transfer (IGT) events, subsequent genomic rearrangements, or the segregation of NUMT-containing chromosomal regions during strain crossing. Consequently, future research should systematically characterize these NUMT loci using high-quality, strain-specific reference nuclear genome assemblies and annotations. Establishing a precise inventory of nuclear-integrated mitochondrial sequences is essential to prevent off-target amplification and improve the accuracy of nuclear and mitochondrial genetic analyses.

## Materials and Methods

### Strain preparation, culture conditions and growth measurements

*Chlamydomonas reinhardtii* strains CC-5325 (*mt^−^*), CC-4425 (*mt^+^*), CC-5816 (*mt^+^*), and CC-125 (*mt^+^*) were obtained from the Chlamydomonas Resource Center (https://www.chlamycollection.org). The remaining CC-5325 lineage strains—CC-124 (*mt^−^*), CC-4051 (*mt^+^*), 17D*−* (*mt^−^*), and CC-4603 (*mt^−^*)— and the field isolated strains (*SI Appendix*, Table S5)—CC-1952 (*mt^−^*), CC-5823 (*mt^−^*), CC-5824 (*mt^+^*), along with CC-5825 (*mt^+^*)—were provided by the Merchant Lab (University of California, Berkeley). To assess bacterial contamination, all strains were initially screened for heterotrophic bacterial contamination on R2A agar plates. Upon confirming the absence of bacterial colonies, seed cultures were grown photoautotrophically in Sueoka’s High Salt Medium (HSM) in 250 mL column bioreactors. Cultures were maintained under continuous illumination at a light intensity of 50 ± 5 µmol photons m^−2^ s^−1^ and sparged continuously with filtered air enriched to 10 % (v/v) CO_2_. Growth was monitored by cell counting using a hemocytometer.

### High-molecular-weight DNA (HMW DNA) extraction and quantification

Cells were collected by centrifugation three days after inoculation (stationary phase) in photobioreactors, yielding approximately 7.5×10^8^ cells per extraction. To minimize DNA fragmentation and preserve mtDNA integrity, a modified HMW DNA extraction protocol was implemented (52). Cell pellets were resuspended in CTAB lysis buffer containing 2% (w/v) cetyltrimethylammonium bromide, 1% (w/v) polyvinylpyrrolidone (molecular weight at 360 kDa), 50 mM Tris-HCl, 20 mM EDTA, and 1.4 M NaCl. The lysis mixture was supplemented with Proteinase K and RNase A (Thermo Fisher Scientific, Waltham, MA) and incubated in a water bath at 65°C for 30 min. Following centrifugation, the supernatant was recovered and gently mixed with phenol: chloroform: isoamyl alcohol (25:24:1, v/v/v). The aqueous phase was collected, treated with a second round of Proteinase K and RNase A for 20 min at 50°C, and subsequently extracted with chloroform: isoamyl alcohol (24:1, v/v). DNA was precipitated from the final recovered supernatant by adding an equal volume of ice-cold isopropanol. The resulting DNA pellet was washed with 70% (v/v) ethanol, air-dried, and resuspend in nuclease-free water overnight. For quality control, DNA purity was assessed using a NanoDrop 2000c spectrophotometer (Thermo Fisher Scientific). DNA concentration was determined using the Qubit 1X dsDNA HS Assay Kit with a Qubit Flex Fluorometer (Thermo Fisher Scientific) according to the manufacturer’s protocols.

### Mitogenomes assembly and annotation of CC-5325

In parallel with the chromosomal assembly of CC-5325, we used HiFi reads to assemble the config 1 mitogenome through a two-step process. First, *de novo* assembly was performed on the complete HiFi read dataset using Flye v2.9.6-b1802 (--meta) (25). Contigs 13*−*20 kb in length with exceptionally high coverage were selected as mitogenome candidates. Collinearity analysis was then conducted between these candidate contigs and the reference mitogenomes (CC-503 and NC_001638.1) using MUMmer v4.0.1 (nucmer --mum) (53). The candidate contig exhibiting high collinearity with the reference sequences was designated as the CC-5325 mitogenome assembly. Because this *de novo* assembly was structurally distinct from the references, we performed a secondary reference-guided assembly for validation. HiFi reads were mapped to the reference genomes using minimap2 v2.30-r1287 (54), and the mapped reads were re-assembled with Flye (--genome-size 30k --asm-coverage 80). This analysis recovered a single contig identical to the initial *de novo* assembly, thereby supporting the novel config 1 mitogenome structure.

To recover the config 2 mitogenome, we filtered the HiFi reads against the config 1 sequence and assembled the retained reads using Flye (--genome-size 16k --asm-coverage 50). This assembly produced a single contig comparable in length to the reference mitogenomes, with sequence collinearity independently confirmed using MUMmer. Finally, HiFi reads were mapped back to both mitogenome using minimap2, sorted with SAMtools v1.21 (55), and visualized in the Integrative Genomics Viewer (IGV v2.19.7) (56) to evaluate assembly quality and structural variants.

To further assess the accuracy and structural differences of the assembled mitogenome, we performed variant calling between the newly assembled config 1 and five published reference mitogenome assemblies: *CW92* (NC_001638.1), *CW15* (U03843.1), CC-124 (EU306622.1), CC-503 (9), and CC-4532 (57). SNPs and INDELs were identified using nucmer and dnadiff functions in MUMmer. For this analysis, the published mitogenomes were designated as references, and the newly assembled config 1 was used as the query baseline.

For mitogenome annotation, we used a hybrid, reference-guided approach to account for the highly conserved nature of mitochondrial genes. Config 1 was initially submitted to GeSeq (58) for draft annotation using NC_001638.1 as reference. Liftoff v1.6.3 (59) was utilized to transfer annotations from the CC-503 reference mitogenome to config 1. These two annotation datasets were then integrated and cross-validated, and redundant GeSeq features were removed. Translated coding sequences from both sources were compared; in cases where the Liftoff annotation conflicted with the GeSeq output, the GeSeq annotation was retained as it directly reflected the config 1 sequence. Gene boundaries were manually verified by visualizing the features in JBrowse 2 (60). Subsequently, tRNAs were re-annotated using tRNAscan-SE v2.0.12 (-O --max) (61), and the resulting annotation replaced the draft tRNA annotations. Finally, the complete annotation was manually reviewed in JBrowse 2 to ensure accuracy.

### Draft genome assembly of CC-5325 lineage strains and evaluation of organelle sequence homology

Whole-genome sequencing (WGS) datasets for the CC-5325 lineage strains, including CC-4603 (SRR1797971), CC-4425 (SRR1797954), and CC-4501 (SRR1797967), and their ancestral strains CC-124 (SRR1797946) and CC-125 (SRR1793656), were retrieved from the NCBI Sequence Read Archive (SRA). Initial read quality was evaluated using FastQC v0.12.1 (https://www.bioinformatics.babraham.ac.uk/projects/fastqc/), and adapter sequences along with low-quality bases were trimmed using Trimmomatic v0.39 (62). A secondary round of FastQC was performed to verify post-trimming quality. *De novo* genome assemblies were generated for each strain using SPAdes v4.1.0 (63) with default parameters. To avoid the loss of unique genomic contigs or misassembled artifacts, the assemblies were retained at the contig level. Nuclear contigs were identified by mapping the assembled contigs to the reference CC-5325 nuclear genome, and contigs classified as organellar or putative contaminant sequences were removed. Redundant duplicated haplotigs within the nuclear assemblies were subsequently removed using purge_dups v1.2.6 (64). To identify sequence homologies between the organelle and nuclear genomes, local alignments were performed using BLASTN with a modified word size (-word_size 6) (65). The config 1 mitogenome sequence was used as the subject database, and curated nuclear contigs from each lineage strain were queried against it separately.

### Read coverage estimation and digital PCR (dPCR) quantification of mitogenome configurations

Unfiltered WGS reads from the strains associated with the CC-5325 lineage and its ancestral strains were independently mapped to the CC-5325 config 1 mitogenome assembly using BWA-MEM (66). The resulting alignments were sorted using SAMtools, and read coverage was visualized in the IGV. A significant reduction in read coverage, accompanied by extensive SNP and INDEL enrichments, was observed near config 1-specific boundary region (∼15,048 bp) (Fig. 1*B*). Because sequence divergence at this region compromised accurate mapping and reads alignment for non-CC-5325 strains, we conducted strain-specific config 1 pseudo-references to systematically account for these SNPs and short INDELs using BCFtools (mpileup and consensus) (55). Raw WGS reads were subsequently re-mapped to their respective pseudo-references. To quantify the relative abundance of the mitogenome configurations, reads with exact-match alignments and no clipping reads across the site 4,001 (left junction) and site 15,048 (right junction) were extracted using SAMtools (view -F 2304). The relative abundance of config 1 within the total mitogenome pool (config 1 + config 2) was calculated as the ratio of reads spanning the right junction to those spanning the left junction.

Because obtaining mtDNA free of nuclear DNA contamination is challenging in *C. reinhardtii*, we used total DNA, including nuclear, mitochondrial and plastid DNA, as the dPCR template. To minimize the interference from NUMTs, we designed three specific primer sets—Link1, Link2, and a BamHI Indicator (BI)—guided by the NUMT distributions across the tested strains (Fig. 4). Such design avoids off-target amplification and any chimeric PCR products. The Link1 primer set targets a region shared by both config 1 and config 2 and was used to estimate the total mitogenome pool. The Link2 primer set targets a config 1-specific region, enabling quantification of this configuration. The BI primer set was included as a positive control and is expected to match the Link1 copy number.

Reactions were prepared in a total volume of 15 µL using QIAcuity EvaGreen PCR Master Mix (QIAGEN, Hilden, Germany) and loaded into QIAcuity 26k 8-well nanoplates (QIAGEN). Total DNA input was optimized to 50 pg per reaction to avoid partition oversaturation and ensure accurate Poisson distribution statistics. The nanoplates were processed using the QIAcuity Digital PCR System (QIAGEN) under the following conditions: initial denaturation at 95 °C for 2 min, followed by 40 cycles of 95 °C for 15 s and 60 °C for 30 s, and a final extension step at 35°C for 5 min. Following thermal cycling, the nanoplates were imaged in the green fluorescence channel with an exposure duration of 200 ms. Absolute copy numbers and data analysis were performed with four independent replicates using the QIAcuity Software Suite (QIAGEN). The relative abundance of config 1 within the total pool was calculated as [config 1 / (config 1 + config 2)].

Because the Link2 target region exhibits structural variation in strains of the CC-5325 lineage, except CC-5325 itself (Fig. 1*B*), the Link2 primer set was not suitable for quantifying mitogenome config 1 in other *Chlamydomonas* strains. For strains other than CC-5325, we used an alternative strategy based on *cob*, Link1, and BI primer sets to determine the config 1 to config 2 ratio (*SI Appendix*, Fig. S8). In dPCR, physically linked target sequences, such as tandem arrays present in the mitochondrial concatemeric structures, frequently co-segregate into a single partition, detected as one positive fluorescent event rather than discrete individual copies. To resolve this linkage-associated artifact, total DNA from strains other than CC-5325 was digested with FastDigest BamHI (Thermo Fisher Scientific) for 20 min prior to dPCR. This BamHI treatment was intended to physically separate the inverted repeats of config 1, thereby increasing the abundance of quantifiable *cob* amplicons (Δ *cob*) while reducing the abundance of intact BI amplicons (Δ BI).

According to the proposed interpretation of the dPCR assay, the increase in detectable *cob* signal following BamHI digestion (Δ *cob*) reflects the release of physically linked *cob* copies associated with config 1-containing mtDNA molecules (config 1 monomers and config 1-containing concatemers), whereas the change in Link1 (Δ Link1) provides an estimate of the contribution of concatemeric mtDNA junctions. Furthermore, the reduction in intact BI amplicons (Δ BI) reflects BamHI cleavage of BI-containing sequences associated with both config 1 and 2, including monomeric and concatemeric molecules. Therefore, the relative abundance of config 1 within the total mitogenome pool was estimated using the following equation:

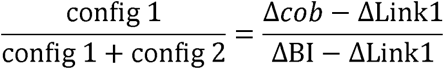

## Data, resources and materials availability

The genomic sequencing datasets used in this study were generated as part of an ongoing telomere-to-telomere (T2T) genome assembly and annotation project for *C. reinhardtii* CC-5325. To ensure reproducibility, the relevant genome assembly and raw reads have been deposited in the NCBI BioProject PRJNA1549149. These publicly available resources include the T2T genome assembly (GenBank accession:), the PacBio HiFi reads (SRA accession: SRR41325606), and the Oxford Nanopore Technologies (ONT) reads (SRA accession: SRR41328809). All other data supporting the findings are available within the article and its Supplementary Information. The bioinformatic software and parameters are detailed in the Materials and Methods, and additional custom code and computational scripts used for this manuscript are publicly available through GitHub at https://github.com/ZhongshiHe/Supplementary-code-source-for-mitochondrial-heteroplasmy-of-Chlamydomonas-reinhardtii.

## Acknowledgments

We gratefully acknowledge Dr. Yibing Zeng (Department of Genetics, University of Georgia) for assistance with genome sequencing and quality assessments. We also thank Dr. Yi-Ying Lee (Institute of Marine and Environmental Technology (IMET), University of Maryland Center for Environmental Science (UMCES)) for assistance with microalgal growth and maintenance. Additionally, we thank the BioAnalytical Services Laboratory at IMET for the provision of the ONT sequencing and dPCR analysis platform. Further appreciation is expressed to Dr. Lauren E. Jonas (Department of Plant Sciences and Plant Pathology, Montana State University) and Dr. Yotam Blech-Hermoni (QIAGEN) for their technical guidance regarding digital PCR operations. This work was supported in part by the DOE Office of Fossil Energy and Carbon Management (FE-0032188) and IMET Angel Investment Fund to Y.L. Funding was also provided by the UMCES Presidential Bridge Funds to Z.H.

## Author Contributions

*Z.H.* and *Y.L.* designed the research, *Z.H.* performed the experiments, analyzed the data and drafted the manuscript; *T.R.B.* independently validated the key analysis; *Y.L.* and *S.D.G.* provided resources; and *Y.L.*, *T.R.B.*, and *S.D.G.* supervised the research progress.

## Competing Interest Statement

The authors have no conflicts of interest to declare.

## Notes

### Competing Interest Statement

The authors have declared no competing interest.

