## Supplementary Information for "Two genomes in one mitochondrion: mitochondrial heteroplasmy and mitogenome dynamics in *Chlamydomonas reinhardtii*"

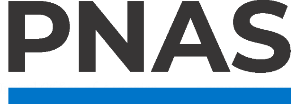

**Supporting Information for**

Two genomes in one mitochondrion: mitochondrial heteroplasmy and mitogenome dynamics in *Chlamydomonas reinhardtii*

Zhongshi He, Tsvetan R. Bachvaroff, Sean D. Gallaher, Yantao Li

Yantao Li

**This PDF file includes:**

Supporting text

Figures S1 to S7

Tables S1 to S5

Legends for Datasets S1 to S7

SI References

**Other supporting materials for this manuscript include the following:**

Datasets S1 to S7

Supporting Information Text

**Supplementary results and discussion**

**"Telomeres" are not present in mitogenomes of *C. reinhardtii* CC-5325 lineage.** Previously, the linear mitogenome of *C. reinhardtii* has been reported to contain terminal "telomeric" structures, consisting of 531-532 bp inverted repeats at both ends that were proposed to mediate mitogenome replication (1). To reexamine these terminal sequences, we mapped our high-accuracy PacBio HiFi reads against the commonly used reference mitogenome (NC_001638.1). The conserved core genomic region exhibited high coverage (> 9,000×) and strong sequence fidelity, with only 24 SNPs, 8 short insertions (3–5 bp), and 1 single-base deletion (Fig. S4*A*). This high degree of sequence conservation validates the mapping quality and is consistent with the evolutionary conservation of the core coding genes of the mitogenome. However, the mapped reads across the terminal regions revealed structural discrepancies that challenged the existence of the proposed "telomeres." Rather than displaying uniformed coverage across the two 532-bp terminal regions, the 5′ (left) terminus exhibited an unusual read depth spike, whereas the 3′ (right) terminus completely lacked reads support for the reference arrangement. Furthermore, sequence analysis confirmed that neither terminal region contained canonical telomeric sequences of Chlamydomonas (e.g., 5′-TTTTAGGG-3′ or 5′-CCCTAAAA-3′), nor did they exhibit any other tandem repeat arrays characteristic of canonical telomeres.

Because the previously proposed telomeric structure was supported in part by targeted PCR analysis, we re-examined the previously reported 0.8-kb PCR product (GenBank: M83998.1), which had been interpreted as evidence for a fused mitogenome terminus (2). Mapping this sequence to the assembled and annotated config 1 mitogenome (Fig. S4*B*) revealed that, although the reported sequence itself was accurate, its interpretation as evidence for native telomeres needs reconsideration. The sequence contained a partial *rrnL2a* and complete *rrnL2b* sequence, followed by the 3′ intergenic region, a 12-bp motif (position 370-381 bp), the complete left terminal region, and a partial *cob* sequence. Importantly, this 12-bp motif presented between the 3′ of the intergenic region and the beginning of the 5′ terminal region, providing a potential site for template switching during PCR amplification. One possible explanation is that, upon reaching the end of a template with the 12-bp motif, the exhaustion of the template likely caused the polymerase to pause and dissociate. Such a partially extended DNA strand (newly synthesized *rrnL2a*/*b*) containing the 12-bp motif may then anneal to the homologous 12-bp motif at the 5′ head of another template (left terminal region) and resume replication (Fig. S4*C*). This template-switching event could generate an artificial fusion of the right core region and the left terminal region. Although this jumping PCR artifact provides a reasonable mechanistic explanation for the previously reported experimental results (2), we hypothesize that this end-to-head junction may actually represent a genuine DNA arrangement within the native mitochondrial DNA population.

**Re-evaluation of historical mitogenome structural artifacts.** In this research, we reevaluated the terminal sequences reported by Ma et al. (2), suggesting that their inverse PCR products may have originated from PCR-mediated template-switching artifacts or authentic junctions on [config 1—config 2] concatemers (Fig. S4*B* and *C*). Another benchmark study supporting the telomere-mitogenome model sought to directly sequence the native DNA ends using the biochemical and sequencing techniques available at the time (1). To accommodate the requirements of plasmid vector ligation, they modified the native DNA ends via enzymatic blunting (employing nuclease S1 for the removal of single-stranded overhangs alongside Klenow fragment-mediated fill-in synthesis) before analyzing the cloned fragments by Sanger sequencing (3). To characterize the native terminal sequences, they analyzed uncloned, restriction-digested fragments by 5′-radioactive labeling (^32^P) followed by Maxam-Gilbert chemical sequencing (4). This analysis revealed identical 3′ single-stranded extensions of 39–41 nucleotides at both mtDNA termini.

Alignment of these historical reference sequences to our config 1 assembly revealed that previously reported terminal sequences correspond to regions with different structural contexts in our mitochondrial genome configurations (Table S3). The left terminal arm sequence (EMBL-EBI Accession: X66484) aligned to an internal intergenic region of config 1, which corresponds to the right terminal region of config 2. The alignment encompassed a partial 3′ *rrnL2b* locus and a 5′ nad5-fragment and extended across nearly the entire left terminal region, leaving only 5 bp unaligned. Because Maxam-Gilbert sequencing bypasses PCR amplification, this structural discrepancy cannot be attributed to PCR-mediated template switching. Instead, the apparent terminal arrangement may reflect DNA fragmentation during nucleic acid isolation or restriction digestion, potentially involving the disruption of the fragile junctions between config 2 and config 1 concatemers. Although the resulting left-arm sequence represented a hybrid arrangement, it successfully captured the left terminal region of the mitogenome. By contrast, the historical right-arm sequence (Accession: X66485) contained an unaligned segment from positions 1,580—1,720 bp, but otherwise aligned to an internal region of config 1 extending from *rrnS3* to a truncated nad5-fragment.

Using the sequencing methodologies available at the time, Vahrenholz et al. identified these sequences as long inverted repeats and reported the absence of canonical tandem telomeric arrays (5). As they interpreted these sequences as definitive, non-complementary physical termini, their model did not favor circularization or concatemer formation as replication intermediates. Nevertheless, subsequent descriptions of the *C. reinhardtii* mitogenome have referred to these terminal inverted repeats as telomeric structures. By reexamining these historical sequence data in conjunction with our *de novo* long-read assembly and physical mapping analysis, we demonstrate that these previously characterized terminal-repeat structures are unlikely telomeres, but rather the internal concatemeric junctions of structurally heterogeneous mtDNA molecules that were dismissed by early models. Collectively, these findings challenge the conventional interpretation of the historical sequences and support an alternative model of mitochondrial genome organization in *C. reinhardtii*.

Beyond the terminal inverted repeats, Vahrenholz et al. also identified 41-nt 3′ single-stranded extensions at both historically defined mtDNA termini (5′-CAACCAAACCGATACTGTTTGCGCACTTAGGCATGCGTAGT-3′), which provided a basis for their proposed linear mtDNA replication model (1). However, alignment of this sequence to our config 1 assembly revealed only a single perfect match, located internally at positions 14,905—14,945 bp on the negative strand. This internal localization is inconsistent with the proposed presence of identical terminal extensions at both ends of mtDNA, questioning the application of their classic linear replication model. Notably, Vahrenholz et al. reported difficulties cloning these fragments using conventional plasmid ligation methods, a limitation driven by the strict vector requirement for blunt or highly specific cohesive termini (1, 6). Although the precise cause of these cloning difficulties remains uncertain, these observations are consistent with our proposed replication-dependent recombination (RDR) model, in which the recovered fragments may have originated from structurally complex mtDNA molecules, including replication intermediates, rather than representing telomeric termini (Fig. 2*B*).

Supplementary Figures

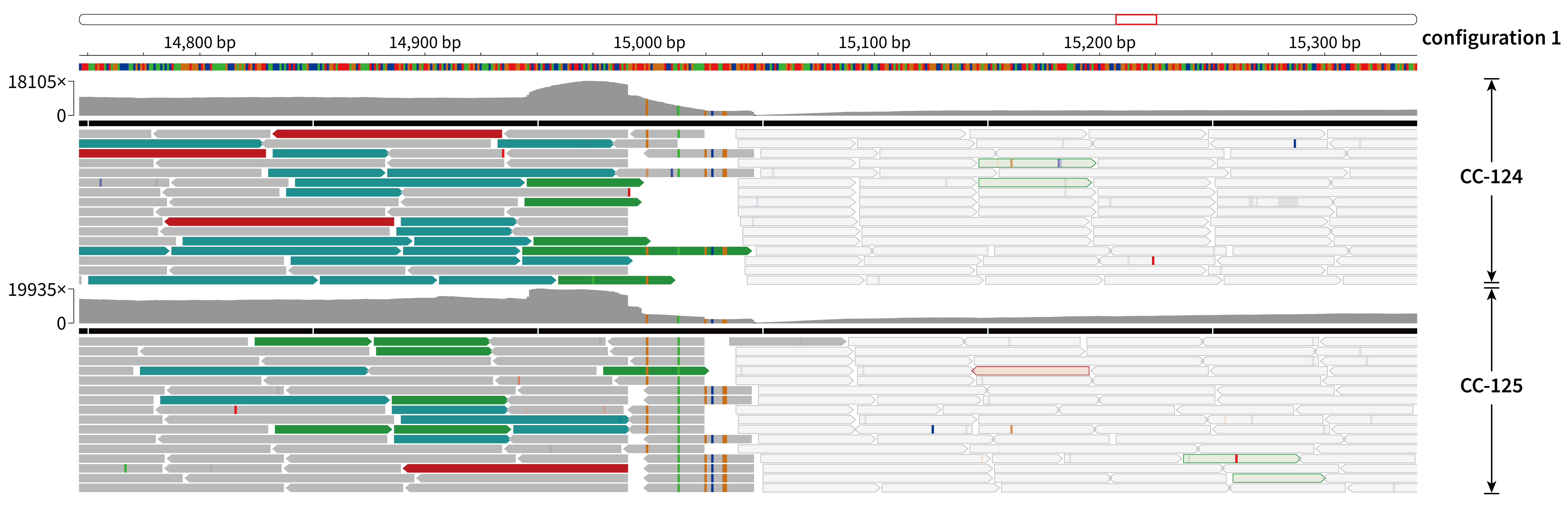

Fig. S1. Whole-genome sequencing read-mapping profiles of *C. reinhardtii* strains CC-124 and CC-125. Illumina paired-end reads from CC-124 (SRR1797946) and CC-125 (SRR1793656) were aligned to the config 1 mitogenome of *C. reinhardtii* CC-5325. The visualized genomic region spans positions 14,750—15,350 bp. Read alignments and coverage depth (upper gray histograms) were displayed using the Integrative Genomics Viewer (IGV) (7). Read colors denote paired-end mapping status, insert size anomalies, and read orientations according to standard IGV color-coding parameters.

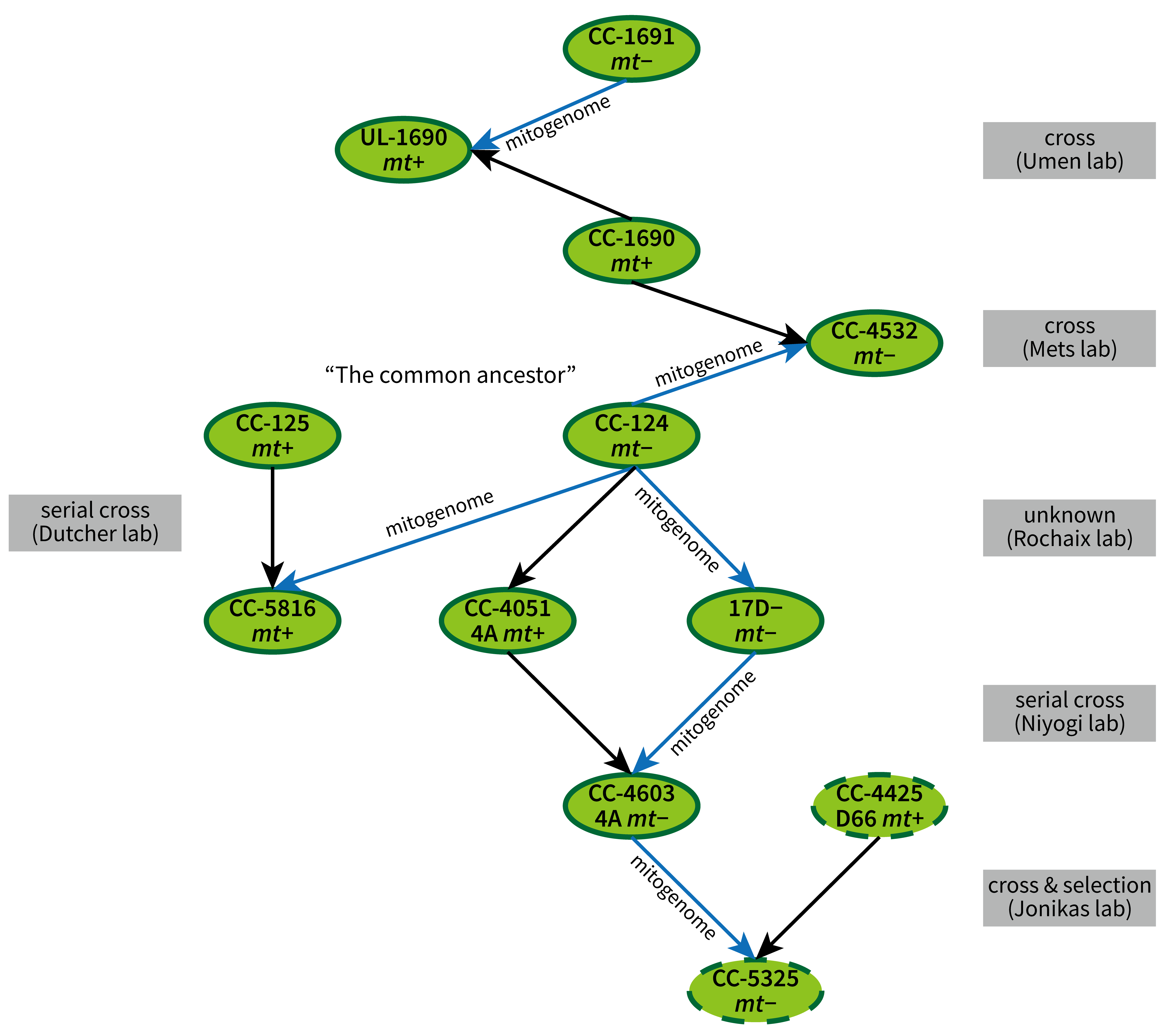

Fig. S2. Strain history and genetic relationships of key *C. reinhardtii* laboratory strains derived from common ancestral isolates (CC-124 and CC-125). Blue arrows indicate the direction of mitogenome inheritance. Gray boxes denote the responsible laboratories, crossing strategies, or selection methods associated with individual lineage transitions. Dashed green borders indicate cell wall-deficient mutants, including CC-4425 and CC-5325. The widely used reference strain, CC-4532 (*mt^−^*) was derived from a cross between CC-124 (*mt^−^*) and CC-1690 (*mt^+^*). CC-5816 (*mt^+^*), for which an additional high-quality genome assembly is available, was isolated following a backcross between CC-125 (*mt^+^*) and CC-124 (*mt^−^*). The laboratory-maintained strain UL-1690 was derived from a cross between CC-1691 (*mt^−^*) and CC-1690 (*mt^+^*).

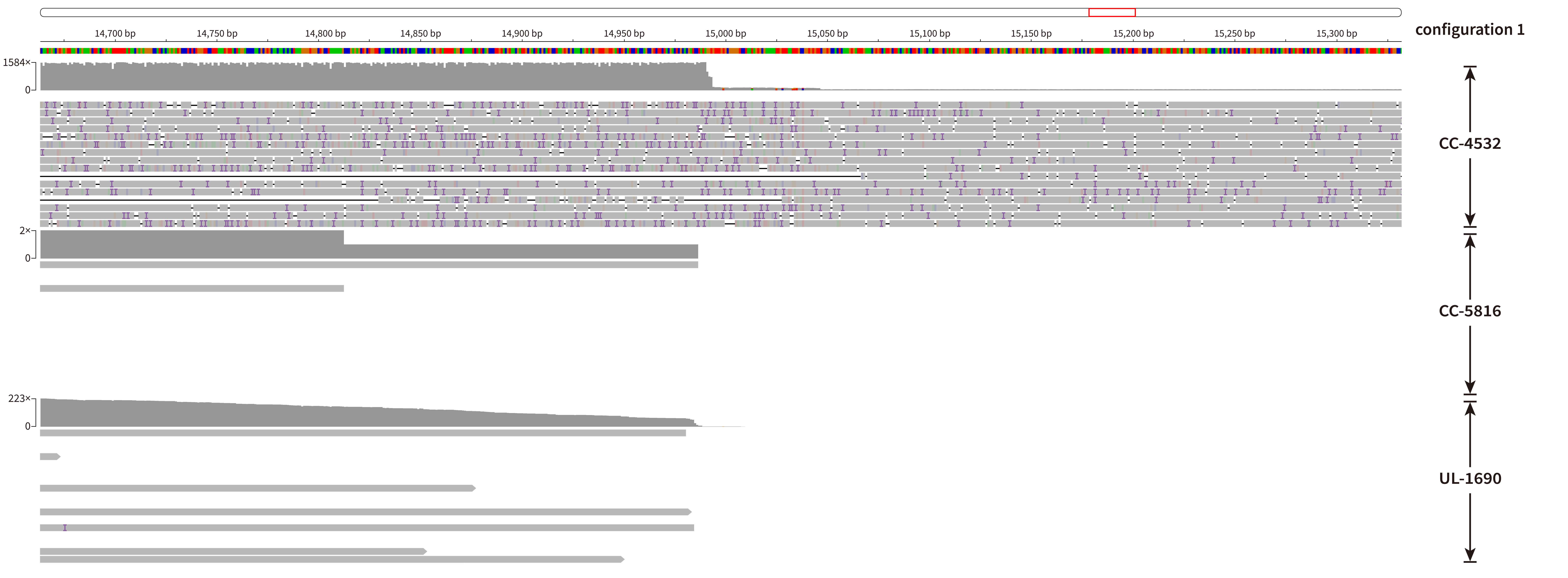

Fig. S3. Long-read whole-genome mapping profiles of *C. reinhardtii* strains CC-4532, CC-5816, and UL-1690. PacBio continuous long reads (CLR) from CC-4532 (BioProject PRJNA887768) and high-fidelity (HiFi) reads from CC-5816 (GenBank: GCA_026108075.1) and UL-1690 (GenBank: GCA_047496495.1) were aligned to the config 1 mitogenome of *C. reinhardtii* CC-5325. The visualized genomic region spans positions 14,650—15,350 bp. Read alignments and coverage depths (gray histograms above the alignments) were displayed using the IGV (7). Individual horizontal gray bars represent aligned single-molecule long reads, and colored vertical marks represent sequence differences relative to the reference genome, including single-nucleotide mismatches, insertions, or deletions, according to standard IGV color-coding parameters.

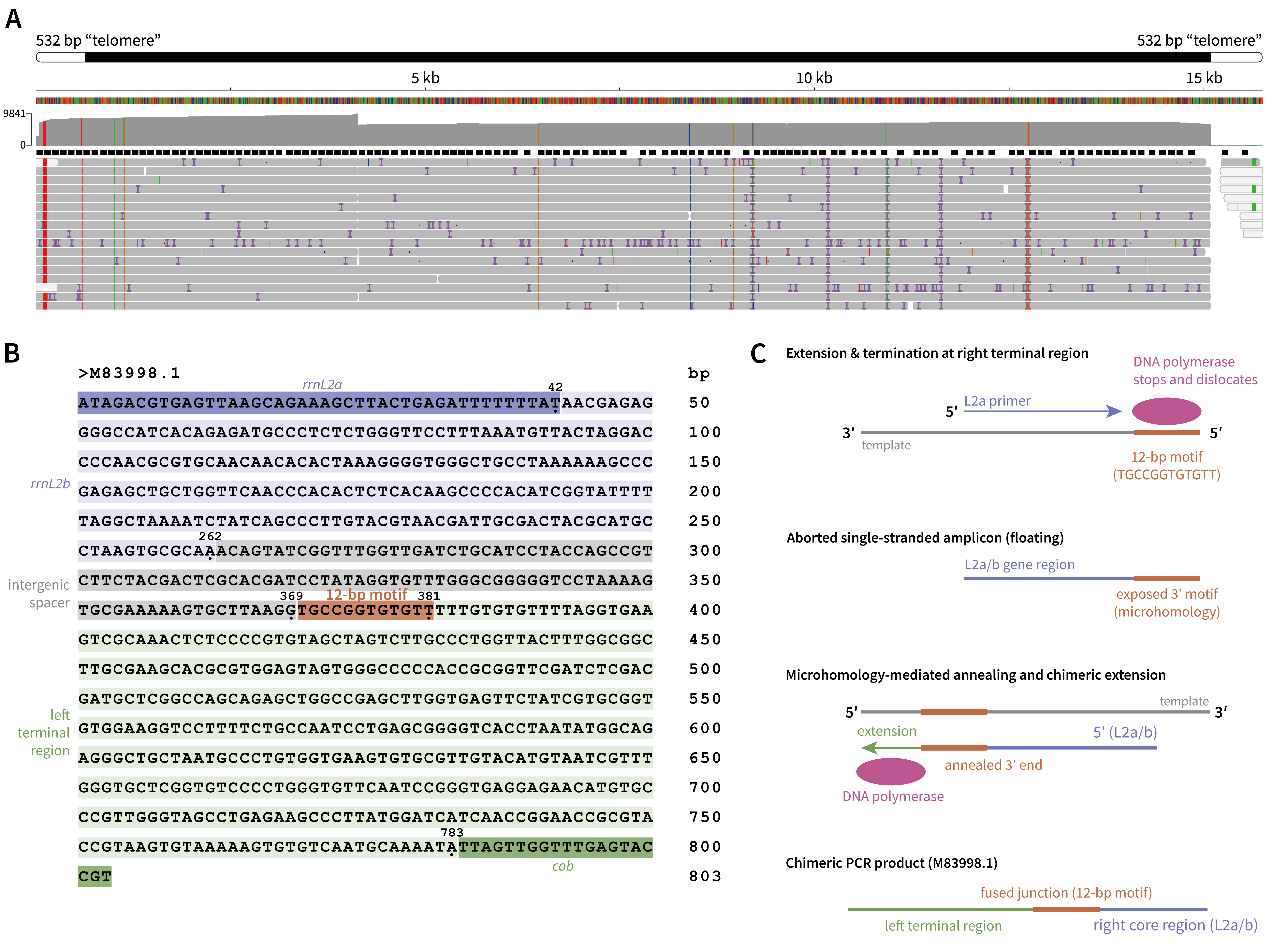

**Fig. S4.** Reinterpretation of historical evidence for terminal telomeres in the *C. reinhardtii* mitogenome. (A) PacBio HiFi reads from CC-5325 were aligned to the NCBI GenBank reference mitogenome (NC_001638.1) and visualized using the IGV (7). The upper tracks mark the position of the historically annotated 532-bp "telomere" region, while the coverage histogram shows reads depth across these regions. (B) Functional re-annotation of the historical 803-bp sequence (GenBank: M83998.1), previously interpreted as structural evidence for linear mitochondrial telomeres. The sequence is reinterpreted as a chimeric fusion joining the right terminal region (purple) to the left terminal region (green), separated by an intergenic spacer (gray) and a shared 12-bp sequence motif (orange). (C) A proposed microhomology-mediated template-switching mechanism during PCR amplification ("jumping PCR") that could generate the chimeric 803-bp sequence (M83998.1).

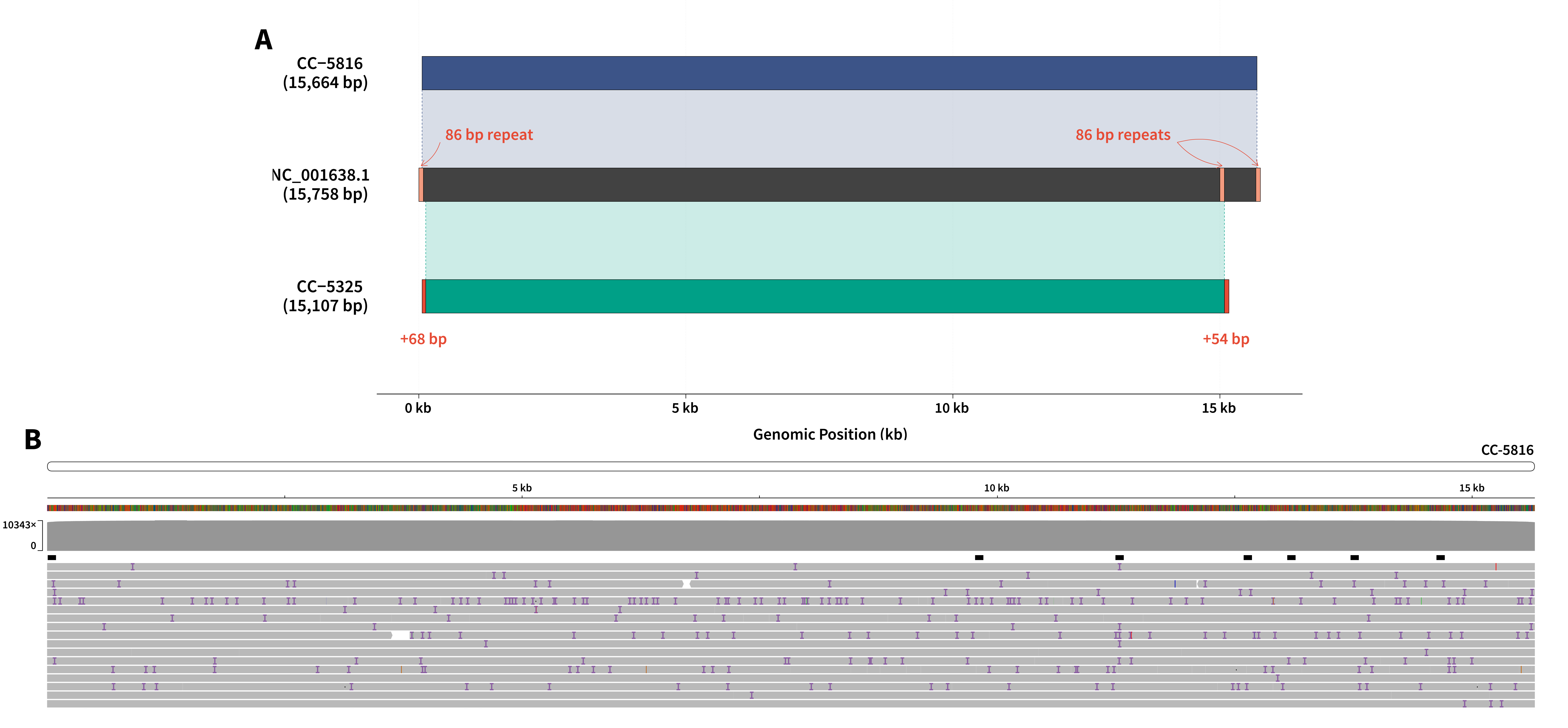

**Fig. S5.** Structural comparison and validation of mitogenome terminal regions in *C. reinhardtii*. (A) Synteny map displays the alignment of mitogenome assemblies from CC-5816 (top) and CC-5325 (bottom) with the reference sequence NC_001638.1 (middle). Colored ribbons represent the highly conserved homologous core regions between the genomes. Vertical dashed lines mark the boundaries of the major aligned core regions. (B) PacBio HiFi reads mapping profile of the CC-5816 mitogenome. The mapped long reads span through the entire mitogenome without soft-clipping in both terminal regions, supporting the sequence continuity and structural consistency of the assembled ends.

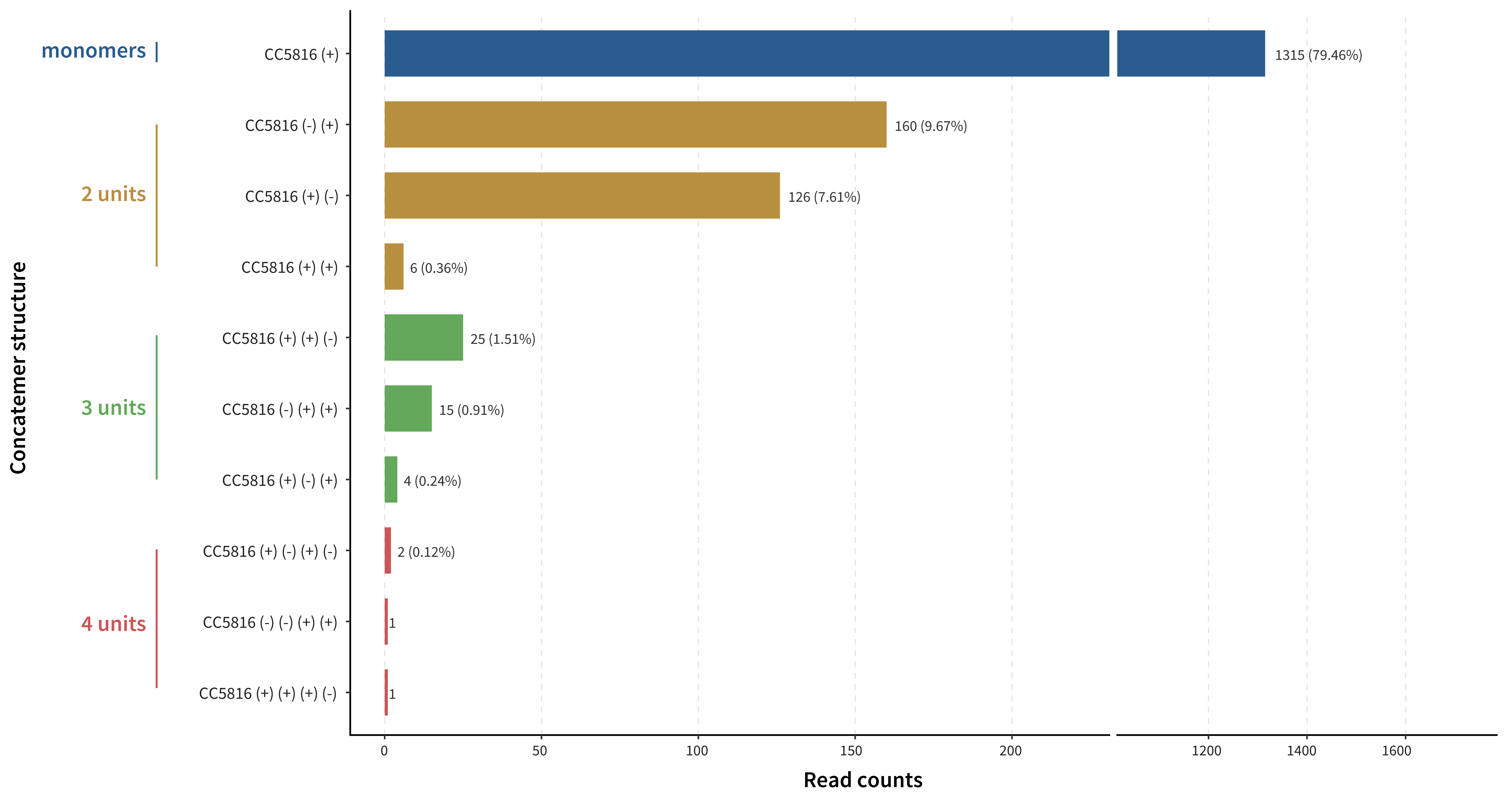

**Fig. S6.** Abundance and structural classification of mitogenome concatemers in *Chlamydomonas reinhardtii* CC-5816. Read counts and relative frequencies (percentages) of ONT long reads corresponding to distinct mitogenome configurations are shown. Structures were classified by the number of mitochondrial genome units (monomers, dimers, trimers, and tetramers) and ordered by abundance within each class. Plus (+) and minus (-) signs denote the relative mapping strand orientations of consecutive mitogenome units. Inverted configurations (e.g., [-] [+] and [+] [-]) constitute the vast majority of multimeric reads, whereas direct head-to-tail concatemers (e.g., [+] [+]) occur at low frequencies. The x-axis breaks discontinuously between 230 and 1,150 reads to accommodate the abundant monomeric class alongside low-abundance multimers.

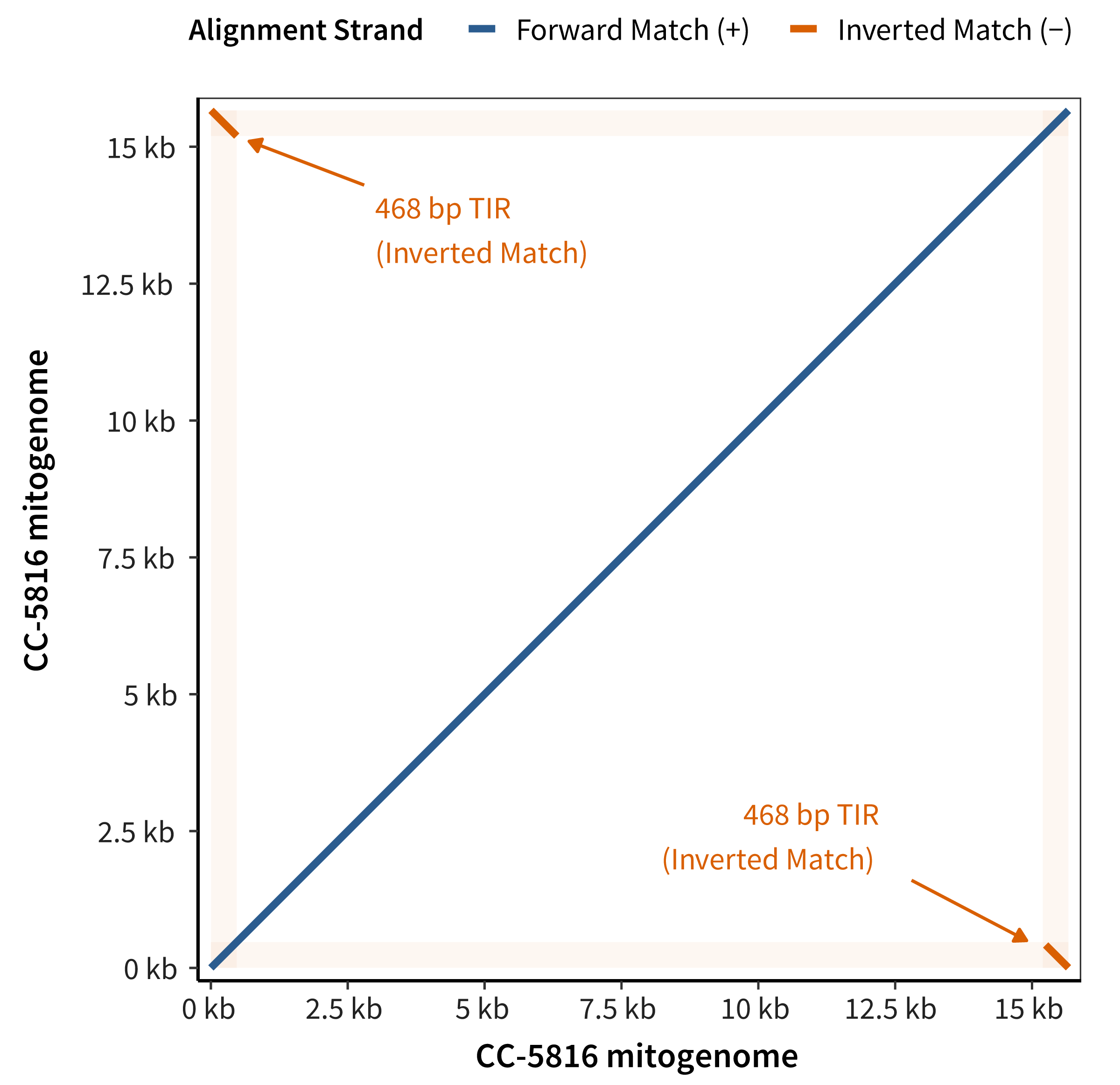

**Fig. S7.** Self-alignment dotplot of the mitochondrial genome of *Chlamydomonas reinhardtii* CC-5816. The self-comparison illustrates sequence collinearity and repeats architecture within the 15,664-bp linear mitogenome. The main diagonal (blue) represents sequence identify along the full-length forward strand (+ strand). The off-diagonal segments at the plot corners (orange) denote 468-bp terminal inverted repeats (TIRs) located at the genome termini (positions 4–471 and 15,197–15,664 bp). Shaded bands along both axes reveal the genomic positions and boundaries of the TIRs. The absence of additional off-diagonal alignments outside the terminal regions suggests that the internal mitogenome sequence lacks extensive repetitive regions.

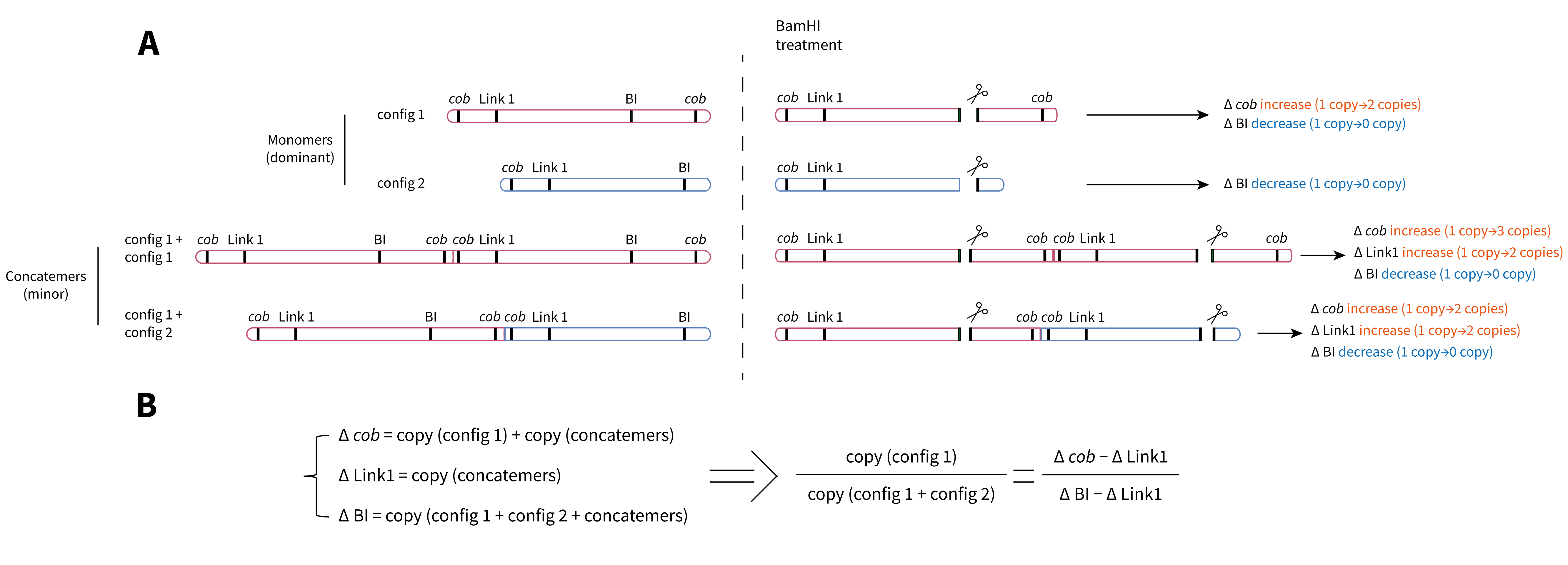

**Fig. S8.** Digital PCR quantification strategy for resolving mitogenome configurations and concatemeric structures. (A) Schematic illustration of a differential dPCR quantification strategy to quantify the dominant monomeric configurations (config 1 and config 2) while accounting for the concatemeric structures. Because physically linked targets may be partitioned together and detected as a single positive partition in undigested DNA, changes in detectable target concentrations (Δ) following BamHI digestion were used to assess the contribution of linked mtDNA segments. BamHI cleavage is expected to produce predictable changes in the concentrations of three amplicons: an increase in *cob* copy number (Δ *cob*) generated by the cleavage of config 1-containing molecules including monomers and concatemers; an increase in Link1 copies (Δ Link1) associated with the separation of mitogenome concatemers; and a decrease in intact BI amplicon copy number (Δ BI), reflecting cleavage of BI-containing sequences associated with the total pool of mitogenomes (config 1, config 2, and their concatemers). (B) Mathematical derivation of config 1 relative abundance. The calculation subtracts the estimated concatemeric contribution (Δ Link1) from both the config 1-associated signal (Δ *cob*) and the total mitogenome pool signal (Δ BI).

Supplementary Tables

Table S1. NUMTs distribution between nuclear genome and mitogenome of CC-5325.

| Homology type | Genome target | Length (bp) | Identity (%) | E-value | Annotation |
| --- | --- | --- | --- | --- | --- |
| NUMT | chr_16 | 42 | 95.238 | 8.94E-08 | Intergenic |
| NUMT | chr_01 | 31 | 100 | 1.33E-05 | Intergenic |
| NUMT | chr_16 | 57 | 94.737 | 2.24E-15 | Intergenic |
| NUMT | chr_17 | 50 | 92 | 1.73E-10 | Intergenic |
| NUMT | chr_01 | 72 | 91.667 | 1.02E-19 | *cob* |
| NUMT | chr_01 | 40 | 97.5 | 7.34E-09 | *cob* |
| NUMT | chr_01 | 333 | 100 | 1.50E-169 | *cob* |
| NUMT | chr_10 | 284 | 88.732 | 8.90E-103 | *nad4* |
| NUMT | chr_03 | 60 | 80 | 0.000162 | *nad4* |
| NUMT | chr_10 | 38 | 92.105 | 4.63E-05 | *nad5* |
| NUMT | chr_16 | 47 | 87.234 | 4.63E-05 | *nad5* |
| NUMT | chr_08 | 37 | 97.297 | 3.12E-07 | *nad5* |
| NUMT | chr_02 | 126 | 95.238 | 4.94E-49 | *nad5* |
| NUMT | chr_09 | 52 | 92.308 | 1.42E-11 | *nad5* |
| NUMT | chr_10 | 63 | 92.063 | 6.43E-16 | *nad5* |
| NUMT | chr_15 | 48 | 97.917 | 3.33E-13 | *nad5* |
| NUMT | chr_02 | 62 | 79.032 | 3.80E-06 | *nad5* |
| NUMT | chr_05 | 77 | 98.701 | 6.02E-29 | *nad5* |
| NUMT | chr_01 | 36 | 100 | 2.56E-08 | *nad5* |
| NUMT | chr_07 | 37 | 97.297 | 3.12E-07 | *cox1* |
| NUMT | chr_02 | 41 | 95.122 | 2.56E-08 | *cox1* |
| NUMT | chr_09 | 280 | 100 | 8.88E-141 | *cox1* |
| NUMT | chr_15 | 384 | 99.479 | 0 | *cox1* |
| NUMT | chr_16 | 72 | 80.556 | 1.09E-06 | *cox1* |
| NUMT | chr_01 | 34 | 100 | 3.12E-07 | *cox1* |
| NUMT | chr_13 | 54 | 98.148 | 2.24E-15 | *cox1* |
| NUMT | chr_17 | 58 | 91.379 | 3.33E-13 | *nad2* |
| NUMT | chr_09 | 62 | 93.548 | 5.28E-17 | *nad2* |
| NUMT | chr_09 | 80 | 81.25 | 7.34E-09 | *nad2* |
| NUMT | chr_07 | 107 | 80.374 | 1.24E-18 | *nad2* |
| NUMT | chr_07 | 62 | 79.032 | 0.000162 | *nad2* |
| NUMT | chr_07 | 57 | 98.246 | 4.33E-18 | *rrnL5* |
| NUMT | chr_02 | 303 | 66.667 | 5.64E-23 | *nad1* |
| NUMT | chr_17 | 33 | 93.939 | 0.000564 | *nad1* |
| NUMT | chr_06 | 156 | 87.821 | 1.72E-48 | *nad1* |
| NUMT | chr_06 | 54 | 90.741 | 4.94E-11 | *nad1* |
| NUMT | chr_09 | 41 | 95.122 | 2.56E-08 | *nad1* |
| NUMT | chr_02 | 29 | 100 | 0.000162 | *nad1* |
| NUMT | chr_10 | 32 | 96.875 | 0.000162 | *rtl* |
| NUMT | chr_01 | 43 | 93.023 | 3.12E-07 | *rtl* |
| NUMT | chr_02 | 56 | 83.929 | 1.09E-06 | *rtl* |
| NUMT | chr_14 | 58 | 96.552 | 1.51E-17 | *rtl* |

Table S1. (*continued*).

| Homology Type | Target Genome | Length (bp) | Identity (%) | E-value | Annotation |
| --- | --- | --- | --- | --- | --- |
| NUMT | chr_07 | 65 | 89.231 | 2.73E-14 | *rtl* |
| NUMT | chr_04 | 49 | 89.796 | 3.12E-07 | *rtl* |
| NUMT | chr_03 | 143 | 98.601 | 1.32E-62 | *rtl* |
| NUMT | chr_02 | 45 | 97.778 | 1.42E-11 | *rrnL8* |
| NUMT | chr_02 | 48 | 93.75 | 6.02E-10 | *rrnL8* |
| NUMT | chr_02 | 43 | 93.023 | 8.94E-08 | *rrnL8* |
| NUMT | chr_08 | 72 | 81.944 | 6.02E-10 | *rrnL8* |
| NUMT | chr_08 | 72 | 81.944 | 6.02E-10 | *rrnL8* |
| NUMT | chr_08 | 72 | 81.944 | 6.02E-10 | *rrnL8* |
| NUMT | chr_01 | 72 | 81.944 | 6.02E-10 | *rrnL8* |
| NUMT | chr_01 | 72 | 81.944 | 6.02E-10 | *rrnL8* |
| NUMT | chr_14 | 72 | 81.944 | 6.02E-10 | *rrnL8* |
| NUMT | chr_14 | 72 | 81.944 | 6.02E-10 | *rrnL8* |
| NUMT | chr_02 | 69 | 81.159 | 3.12E-07 | *rrnL4* |
| NUMT | chr_05 | 33 | 93.939 | 0.000564 | *rrnL1* |
| NUMT | chr_05 | 33 | 93.939 | 0.000564 | *rrnL1* |
| NUMT | chr_12 | 103 | 90.291 | 4.94E-30 | *rrnL2b* |
| NUMT | chr_01 | 44 | 88.636 | 1.33E-05 | *nad5* |
| NUMT | chr_05 | 77 | 98.701 | 6.02E-29 | *nad5* |
| NUMT | chr_02 | 62 | 79.032 | 3.80E-06 | *nad5* |
| NUMT | chr_15 | 48 | 97.917 | 3.33E-13 | *nad5* |
| NUMT | chr_10 | 63 | 92.063 | 6.43E-16 | *nad5* |
| NUMT | chr_09 | 52 | 92.308 | 1.42E-11 | *nad5* |
| NUMT | chr_02 | 126 | 95.238 | 4.94E-49 | *nad5* |
| NUMT | chr_08 | 37 | 97.297 | 3.12E-07 | *nad5* |
| NUMT | chr_16 | 47 | 87.234 | 4.63E-05 | Intergenic |
| NUMT | chr_10 | 38 | 92.105 | 4.63E-05 | Intergenic |
| NUMT | chr_03 | 60 | 80 | 0.000162 | *nad4* |
| NUMT | chr_10 | 284 | 88.732 | 8.90E-103 | *nad4* |
| NUMT | chr_01 | 333 | 100 | 1.50E-169 | *cob* |
| NUMT | chr_01 | 40 | 97.5 | 7.34E-09 | *cob* |
| NUMT | chr_01 | 72 | 91.667 | 1.02E-19 | *cob* |
| NUMT | chr_17 | 50 | 92 | 1.73E-10 | Intergenic |
| NUMT | chr_16 | 57 | 94.737 | 2.24E-15 | Intergenic |
| NUMT | chr_16 | 42 | 95.238 | 8.94E-08 | Intergenic |
| NUMT | chr_01 | 31 | 100 | 1.33E-05 | Intergenic |

Table S2. Genome assembly information of the CC-5325 lineage strains.

| Strains | Mating type | Assembly level | Genome size (bp) | Coverage |
| --- | --- | --- | --- | --- |
| CC-5325 | minus | Telomere-to-telomere | 114,825,738 | 100% |
| CC-4603 | minus | Contig | 88,766,580 | 77.31% |
| CC-4425 | plus | Contig | 88,533,319 | 77.10% |
| CC-4501 | plus | Contig | 90,472,623 | 78.79% |
| CC-124 | minus | Contig | 87,064,442 | 75.82% |

Table S3. Alignment of published *C. reinhardtii* mitochondrial genome fragments to the config 1 assembly.

| Reference accession | Reference coordinate (bp) | Mitogenome config 1 coordinate (bp) | Strand | Annotated features in target region |
| --- | --- | --- | --- | --- |
| X66484 (Left Arm) | 0–142 | 14,904–15,046 | + | Intergenic region (flanking 3' *rrnL2b* and 5' nad5-fragment) |
| X66484 (Left Arm) | 130–7,968 | 5–7,844 | + | *cob*, *nad4*, *nad5*, *cox1*, *nad2*, *nad6* |
| X66485 (Right Arm) | 0–1,579 | 13,411–14,990 | + | *rrnS3*, trnM-CAU, *rrnL4*, *rrnS4*, *rrnL1*, *rrnL2a*, *rrnL2b* |
| X66485 (Right Arm) | 1,721–2,123 | 18,632–19,034 5–407 | + - | Intergenic region |
| X66485 (Right Arm) | 2,111–2,253 | 14,904–15,046 | - | Intergenic region (flanking 3' *rrnL2b* and 5' nad5-fragment) |

Table S4. Sequences of primer sets used in digital PCR system.

| Primer | Sequences |
| --- | --- |
| *cob*_F | ACCGTGGGCAAACTGAGTCTC |
| *cob*_R | GCTATTGACCTGGTTGGGTGGTAA |
| Link1_F | GCACCGAACACACCAGCCATAA |
| Link1_R | CCCAGTATCCGCCCTAATTCACG |
| BI_F | CCAATACGGGCATCGGGCAAA |
| BI_R | CCATCCCAGAACGGCTGTAGCA |
| Link2_F | TCTTCTACGACTCGCACGATCCT |
| Link2_R | CCAGACTAACCATCATATAGCCAAGTTGA |

| Table S5. Information of the field-isolated Chlamydomonas strains. | | | | | |
| --- | --- | --- | --- | --- | --- |
| Strains | Original designation | Mating type | Geographic origin | Collection year | Collector |
| CC-1952 | S1 C5 | minus | Minnesota, USA | 1986 | P.A. Lefebvre |
| CC-5823 | CC-2342 (Jarvik #6) | minus | Pennsylvania, USA | 1989 | Jonathan Jarvik |
| CC-5824 | CC-2343 (Jarvik #224) | plus | Florida, USA | 1989 | Jonathan Jarvik |
| CC-5825 | CC-2936 (LEE-2) | plus | Quebec, Canada | 1993 | Graham Bell |

Dataset S1 (separate file). Mitogenome configuration 1 sequence of *Chlamydomonas reinhardtii* CC-5325 in fasta format.

Dataset S2 (separate file). Mitogenome configuration 1 annotation of *Chlamydomonas reinhardtii* CC-5325 in gff3 format.

Dataset S3 (separate file). Mitogenome configuration 2 sequence of *Chlamydomonas reinhardtii* CC-5325 in fasta format.

Dataset S4 (separate file). Mitogenome configuration 2 annotation of *Chlamydomonas reinhardtii* CC-5325 in gff3 format.

Dataset S5 (separate file). Raw PacBio HiFi and ONT (Oxford Nanopore Technology) reads mapping to mitogenome config 1 and 2 elements of *Chlamydomonas reinhardtii* CC-5325, revealing native conformations and inter-genomic seams. Config 1 and 2 are represented by length, which are 19 kb and 15 kb, respectively.

Dataset S6 (separate file). Mitogenome assembly of *Chlamydomonas reinhardtii* CC-5816 in fasta format

Dataset S7 (separate file). ONT (Oxford Nanopore Technology) reads from *Chlamydomonas reinhardtii* CC-5816 (NCBI BioProject PRJNA840945) mapping to CC-5816 mitogenome, revealing its native concatemer structure.
